# GroEL/ES buffers entropic traps in folding pathway during evolution of a model substrate

**DOI:** 10.1101/2020.05.12.090233

**Authors:** Anwar Sadat, Satyam Tiwari, Kanika Verma, Arjun Ray, Mudassar Ali, Vaibhav Upadhyay, Anupam Singh, Aseem Chaphalkar, Asmita Ghosh, Rahul Chakraborty, Kausik Chakraborty, Koyeli Mapa

## Abstract

The folding landscape of proteins can change during evolution with the accumulation of mutations that may introduce entropic or enthalpic barriers in the protein folding pathway, making it a possible substrate of molecular chaperones *in vivo*. Can the nature of such physical barriers of folding dictate the feasibility of chaperone-assistance? To address this, we have simulated the evolutionary step to chaperone-dependence keeping GroEL/ES as the target chaperone and GFP as a model protein in an unbiased screen. We find that the mutation conferring GroEL/ES dependence *in vivo* and *in vitro* encode an entropic trap in the folding pathway rescued by the chaperonin. Additionally, GroEL/ES can edit the formation of non-native contacts similar to DnaK/J/E machinery. However, this capability is not utilized by the substrates *in vivo*. As a consequence, GroEL/ES caters to buffer mutations that predominantly cause entropic traps, despite possessing the capacity to edit both enthalpic and entropic traps in the folding pathway of the substrate protein.

## INTRODUCTION

Mutations in a protein sequence may subtly change either thermodynamics of the folding polypeptide or protein-solvent interactions. *In vivo*, mutations that arise spontaneously may lead to problems in the folding pathway or stability of proteins. This, in turn, may make the proteins either non-functional or dependent on molecular chaperones for folding (Chiti and Dobson, 2017; Hartl, 2017)

In *E. coli*, the most abundant cytosolic chaperone systems consist of the Hsp70 system (DnaK/DnaJ/GrpE), the Hsp60 system (GroEL/GroES), Hsp90 (HtpG), Trigger Factor (Tig) and SecB along with other less abundant chaperones and small heat shock proteins. Properties of the substrates assisted by these chaperone systems have been explored by multiple groups (Calloni et al., 2012; Dunn et al., 2001; Houry et al., 1999; Kerner et al., 2005; Knoblauch et al., 1999; Rüdiger et al., 1997). While the DnaK system binds to many proteins and has the potential to stabilize thermosensitive proteome (Zhao et al., 2019), GroEL/ES binds and helps in the folding of a much smaller subset of cellular proteins (Kerner et al., 2005; Niwa et al., 2016). The mechanism of substrate targeting to the right chaperone is an active field of research underlining the significance of the conformations adopted by proteins in their non-folded states (Mapa et al., 2012; Nagpal et al., 2015).

While canonical chaperone-targeting is important for chaperone-dependent wild type proteins, the accumulation of mutations during evolution may create additional substrates requiring chaperone assistance. Can we predict the type of mutations on a GroEL/ES independent protein that would make it GroEL/ES dependent? Mechanistically, some mutations may increase the propensity of formation of non-native contacts (intramolecular or intermolecular) that need to melt before the protein can fold to its native state (enthalpic trap) while other mutations may increase the flexibility of the non-native states (entropic trap), both of which are folding problems at opposite ends of the spectrum. Do different chaperone systems differ in their capacity to edit the two types of mutations?

GroEL/ES system is capable of removing the entropic trap in its substrates (Chakraborty et al., 2010; Georgescauld et al., 2014). It has also been shown to unfold proteins (Sharma et al., 2008) and remove enthalpic traps in the folding pathway *in vitro*. Does the chaperonin possess both these activities *in vivo*? When proteins accumulate mutations, which type of mutations would be preferentially accommodated by GroEL/ES? This is difficult to answer with the previous model substrates as neither the authentic substrates (which precludes understanding the mutational steps that led to its chaperone dependence) nor the slow-folding model substrates (that were not evolved for GroEL/ES dependence *in vivo* in an unbiased manner) gave us the handle to quantitate folding assistance by GroEL/ES *in vivo*. Although, Horovitz group made advances in this direction by identifying that mutations in frustrated sequence regions lead to GroEL/ES dependence *in vivo*, but the biophysical consequence of these mutations on the folding landscape was not investigated (Bandyopadhyay et al., 2017).

To address this, we sought an unbiased screen using mutagenesis to obtain an *in vivo* GroEL/ES substrate starting from a spontaneously folding GroEL/ES-independent substrate. We show that the identified substrate is exclusively dependent upon the GroEL/ES system *in vivo* and *in vitro*. We find that the unique mutation present in the pool of GroEL/ES dependent protein resulted in an entropic trap in folding that is corrected by GroEL/ES system. We also show that DnaK/DnaJ/GrpE (KJE) system or the GroEL chaperone can take care of enthalpic traps effectively *in vitro* in an ATP dependent manner. This function of GroEL is essential for folding the substrate protein *in vivo*. Thus, we posit that the proteins that acquire entropic traps during evolution would be assisted by GroEL/ES system *in vivo*. While GroEL/ES also possesses the capability to edit folding landscape by preventing the formation of non-native contacts, this chaperoning capacity is not exclusive to GroEL/ES but is also shared by more abundant KJE system.

## RESULTS

### Isolation of a synthetic GroEL-dependent substrate

A single mutational step may make a spontaneously folding protein chaperone-dependent during evolution. To learn the physico-chemical basis of the mutational step that confers chaperone-dependence, we wanted to mimic this evolutionary step and develop a synthetic substrate dependent on the canonical chaperone GroEL/ES *in vivo* from a GroEL/ES-independent protein. Comparison of the mutant and Wt protein would help in understanding the type of folding problems that GroEL/ES tends to edit, and the mechanism of its chaperoning action. We chose yeGFP (yeast enhanced GFP-referred to as Wt GFP hereafter), a fast-folding form of GFP, as the starting protein as, 1) folding of the Wt protein was independent of GroEL/ES *in vivo* and *in vitro* (Figure 1A), and 2) it was easy to quantify the amount of soluble well-folded protein in single-cells using flow-cytometry (Verma et al., 2020). Importantly, to normalize for expression levels, plasmid copy number and induction, we used Wt mCherry in an operonic construct containing Wt GFP followed by ribosome-binding-site (RBS) and Wt mCherry, under the control of an arabinose-inducible system (Figure 1B). The expression levels of mCherry were used as a reference to obtain the relative expression levels of GFP (Verma et al., 2020).

**Figure 1:**
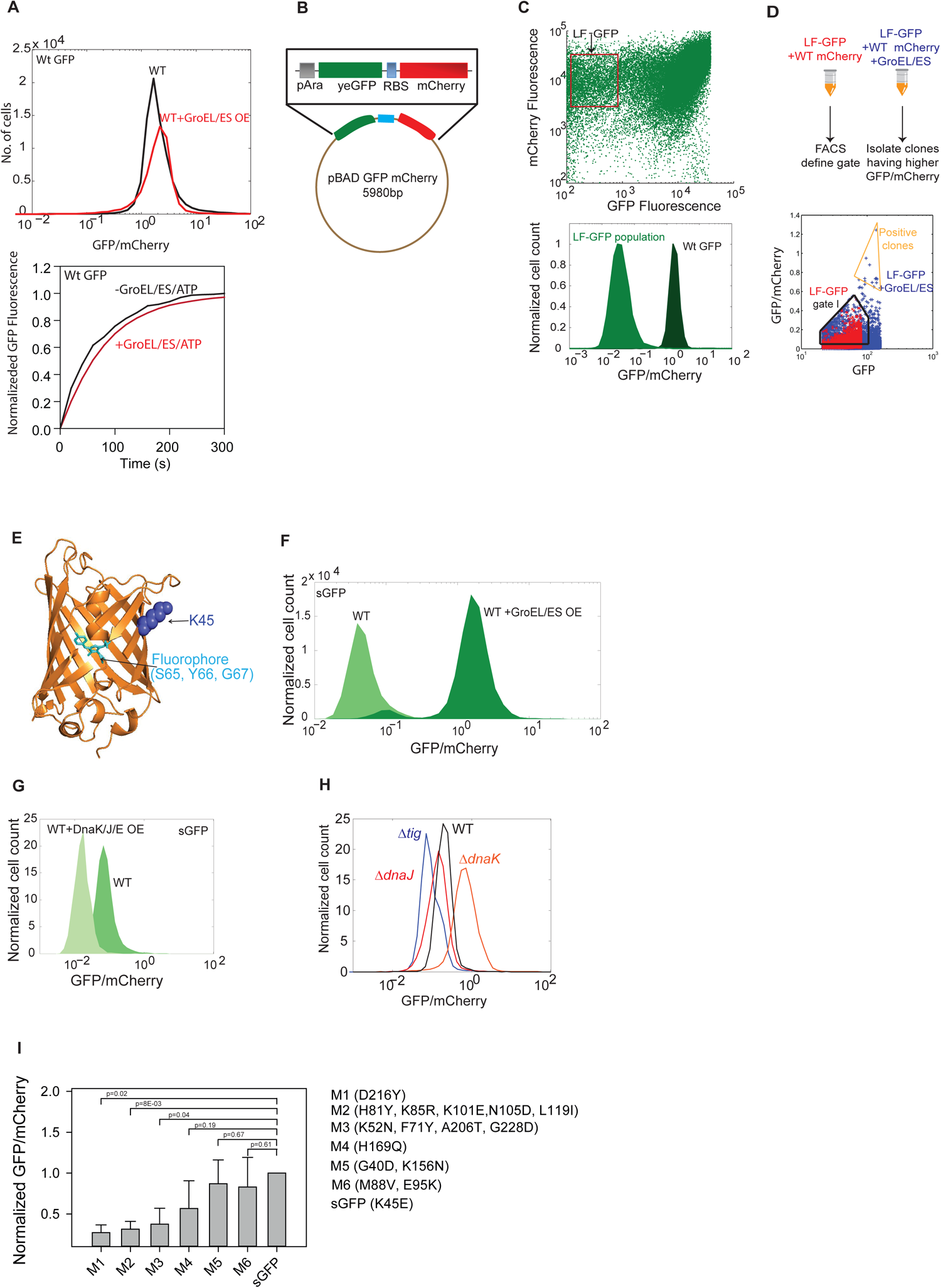
Laboratory evolution of an authentic substrate of GroEL/ES. (A) Upper panel: Histogram for *in vivo* fluorescence of Wt GFP (represented as the ratio of GFP/mCherry) in the presence and absence of plasmid-based overexpression of GroEL/ES. Lower panel: Refolding kinetics of Wt GFP in the presence and absence of GroEL/ES/ATP. Spontaneous refolding of Wt GFP was initiated by a 100-fold dilution of the unfolded proteins in buffer A (20 mM Tris, 20 mM KCl, 5 mM MgCl_2_, 2 mM DTT, pH 7.4) at 25°C. For GroEL/ES assisted refolding unfolded protein was diluted 100-fold in buffer A containing 400 nM of GroEL (all concentrations of GroEL are in terms of tetradecamer), 800 nM GroES (all concentration of GroES are in terms of heptamer) so that the final concentration of unfolded protein is 200 nM. Subsequently, refolding was initiated by adding 2 mM ATP. Recovery of GFP fluorescence over time was followed to monitor refolding. (B) Schematic of the bicistronic construct of GFP and mCherry drove by an Arabinose inducible system. RBS indicates the position of the additional ribosome binding site (RBS) to initiate translation of mCherry. (C) Upper panel: Scatter plot of *E. coli* WT cells expressing random mutant library of GFP. The red box highlights the population of cells with low GFP fluorescence (LF-GFP) compared to Wt GFP, with the same mCherry fluorescence as that of Wt GFP. Lower panel: Histogram for flow cytometry of cells harboring low-fluorescent (LF) mutant pool of GFP and the Wt GFP. GFP/mCherry ratio is monitored at a single-cell level. (D) Upper panel: Schematic of FACS to isolate GroEL/ES-dependent mutant GFP clones. Lower panel: The cells harboring LF-GFP library were induced to express GFP mutants either alone or along with the GroEL/ES system. Negative gating (gate I) was defined by the single-cell GFP/mCherry ratio of the LF-GFP when expressed alone (red dots). Separately, LF-GFP expression was induced in the presence of overexpressed GroEL/ES (blue dots), Positive clones were sorted by gating for cells that had higher GFP/mCherry ratio than Gate I (Positive gate). Gate I included cells only with high GFP fluorescence along with a high GFP/mCherry ratio. (E) K45 residue depicted on the crystal structure of GFP (1GFL (Xu et al., 1997; Yang et al., 1996)) showing its surface exposure away from the buried fluorophore. Some of the residues in beta-sheets facing the viewer are not shown to make the fluorophore underneath clearly visible. Only one chain of GFP is shown for clarity. Depiction made using Chimera (Pettersen et al., 2004). (F) Histogram of single-cell GFP/mCherry ratio of sGFP (GFP (K45E)) in the presence and absence of GroEL/ES over-expression. For individual histograms of GFP and mCherry please see Figure S1C. (G) Histogram of single-cell GFP/mCherry ratio of sGFP in the presence and absence of overexpressed DnaK/J/E. For individual histogram of GFP and mCherry please see Figure S1F. (H) Histogram for in vivo GFP/mCherry fluorescence of sGFP in wild type E.coli K12 (WT) and knockout strains for canonical molecular chaperones dnaK (Δ*dnaK*), dnaJ (Δ*dnaJ*) and trigger factor (Δ*tig*). For individual histogram of GFP and mCherry please see Figure S1G. (I) Bar plot for in vivo GFP/mCherry fluorescence of sGFP and 6 independent GFP mutants isolated from LF-GFP library in wild type *E.coli* K12 (WT) overexpressing GroEL/ES. See Figure S1.

To make a GroEL/ES dependent substrate, we used a random mutant library of GFP, constructed using the backbone of this GFP-mCherry operonic construct (Verma et al., 2020). The mutant library showed populations of cells containing varying degrees of GFP fluorescence starting from very low fluorescence to near-Wt GFP fluorescence (Figure 1C, upper panel). Using Fluorescence Assisted Cell Sorting (FACS) we first purified a population of clones (Population-LF) that exhibit extremely low GFP fluorescence compared to Wt GFP (Figure 1C). These clones would either have problems in folding or have quenched fluorescence due to the alteration of the fluorophore environment. We co-transformed Population-LF in a K-12 strain of *E. coli* (BW35113, referred to as WT hereafter) overexpressing GroEL/ES from a plasmid under the control of an IPTG-inducible *tac* promoter. We isolated mutant clones that exhibited higher GFP fluorescence upon GroEL/ES overexpression (Figure 1D and S1A). GFP-mCherry plasmids were isolated from individual clones after sorting and re-transformed to confirm the dependence of GFP fluorescence on the overexpression levels of GroEL/ES *in vivo*. Ten of the isolated clones were sequenced and all of them were found to have the mutation K45E; the mutation did not map to any residues around the GFP fluorophore (Figure 1E). The purified mutant protein had similar *in vitro* fluorescence as that of Wt GFP (Figure S1B). *In vivo* fluorescence of K45E mutant (hereafter referred to as slow-folding GFP or sGFP) increased upon co-expression of GroEL/ES (Figure 1F and S1C). sGFP showed compromised folding and the nascent chains were rapidly cleared *in vivo* in the absence of GroEL/ES overexpression in WT *E*.*coli* cells (Figure S1D). The dependence of sGFP on GroEL/ES for folding was further confirmed with by the drastically enhanced solubility of the protein along with GroEL/ES overexpression (Figure S1E) in BL21(DE3) cell lacking major protease systems. This corroborated well with the increased fluorescence and indicated that sGFP was a folding mutant of Wt GFP that depends upon the GroEL/ES system for folding, *in vivo*.

Furthermore, to check the folding dependence of sGFP on other chaperone systems, we measured the *in vivo* fluorescence of sGFP in the presence of DnaK/J/E overexpression (Figure 1G and S1F). The fluorescence of sGFP did not increase rather decreased in the presence of DnaK/J/E overexpression probably due to the binding of sGFP to DnaK/J/E routing it for degradation. Additionally, deletion of canonical molecular chaperones *Tig, dnaK*, and *dnaJ* did not significantly decrease sGFP fluorescence suggesting that increase in fluorescence of sGFP was not routed through *Tig, dnaK*, and *dnaJ* (Figure 1H and S1G). In fact, GFP fluorescence mildly increased in Δ*dnaK*, which is known to overexpress GroEL/ES system (McCarty and Walker, 1994; Verma et al., 2020). This suggested that sGFP folding was primarily dependent on the cellular GroEL/ES system. Taken together we found that a single mutation on Wt GFP conferred sGFP with a stringent GroEL/ES dependence *in vivo*.

To check if K45E conferred GroEL/ES dependence specifically or any marginally active mutant of GFP would be GroEL/ES substrate, we picked 10 clones randomly from the LF-GFP mutant pool. Out of these 10 clones 6 clones were finally taken to check their dependence on GroEL/ES for their folding. The basal level fluorescence of these mutants was extremely low similar to that of sGFP. Upon overexpression of GroEL/ES with these mutants only three of the six mutants M4, M5, M6 showed an increase in GFP fluorescence like that of sGFP (Figure 1I and S1H). All the others showed lower fluorescence indicating that mutations that confer GroEL/ES dependence are special and are present in ∼50% of the mutant pool. That GroEL/ES expression affects a subset of the mutant GFP pool, indicating that GroEL/ES most likely rescues only specific problems in folding pathways. K45E mutation captured a rare step that would confer GroEL/ES dependence to a GroEL/ES independent folder such as Wt GFP used in the study.

### Refolding of sGFP is limited by a flexibility dependent kinetic trap

To understand the folding barrier introduced by K45E, we characterized the spontaneous refolding pathway of sGFP. Wt GFP and sGFP were unfolded in 6M GuHCl for one hour, causing complete unfolding of the protein (Figure S2A) and refolded by a hundred-fold dilution of the unfolded protein in refolding buffer. We found that sGFP refolded with a much slower rate than Wt GFP (Figure 2A) and was fit well to single exponential kinetics (Figure S2B) with an apparent rate of ∼0.2×10^−3^s^-1^. Additionally, we found that the refolding rate and amplitude of sGFP at 25°C were independent of the concentration of the protein between 12.5 nM to 400 nM (Figure 2B, S2C, and S2D). Similarly, the rates and yield were independent of protein concentration even at 37°C (Figure 2C) suggesting that the protein refolding was not limited by off-pathway aggregation in the temperatures used for this study. This suggested that the mutation K45E perturbed the unimolecular rate of folding in sGFP.

**Figure 2:**
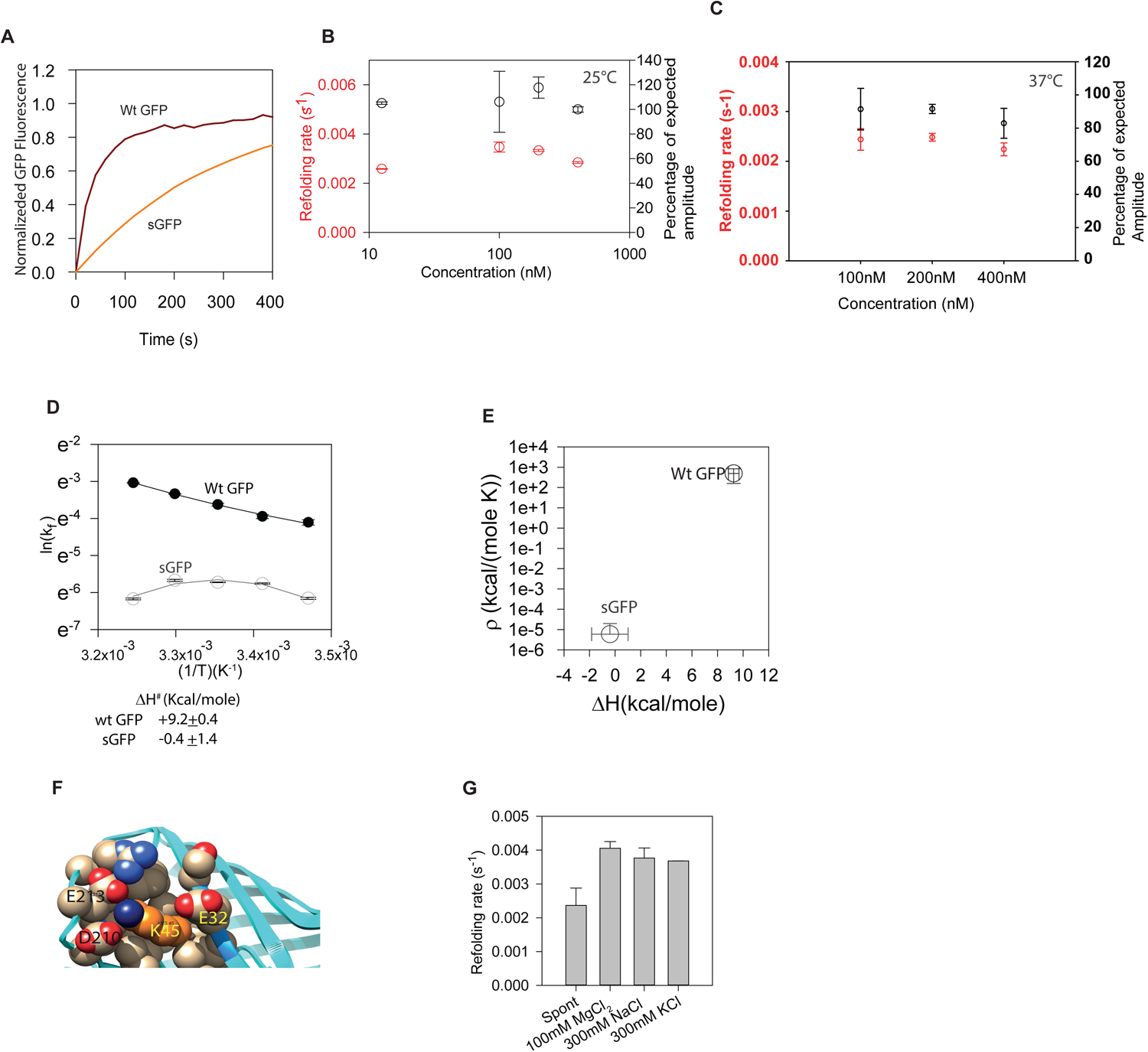
A comprehensive folding landscape for sGFP. (A) Comparison of the refolding kinetics of WtGFP and sGFP. WtGFP and sGFP were unfolded in 6 M GuHCl in buffer A for 1 hour at 25°C. Refolding was initiated by a 100-fold dilution of the unfolded proteins in buffer A (20 mM Tris, 20 mM KCl, 5 mM MgCl_2_, 2 mM DTT, pH 7.4) at 25°C so that the final concentration of the proteins was 200 nM. Refolding was monitored by following the fluorescence of GFP with time. (B) Refolding rate and amplitude as a function of sGFP concentration. sGFP was unfolded as described earlier at 25°C but at different concentrations. Refolding was initiated from each of these by a 100-fold dilution into buffer A at 25°C such that the final protein concentrations were 12.5, 100, 200, or 400 nM. Refolding was monitored by measuring the recovery of GFP fluorescence over time. The refolding traces were fitted to obtain the apparent folding rate (red circles) and the percentage of expected amplitude (black circles). These were plotted as a function of protein concentration. The expected amplitude was calculated based on the fluorescence recovery upon refolding protein at 400 nM concentration. (C) Refolding rate as a function of sGFP concentration. sGFP was unfolded as described earlier at 37°C but at different concentrations. Refolding was initiated from each of these by a 100-fold dilution into buffer A at 37°C such that the final protein concentrations were 100, 200, or 400 nM. Refolding was monitored by measuring the recovery of GFP fluorescence over time. (D) Arrhenius plot for sGFP and Wt GFP refolding. Refolding of sGFP or Wt GFP was initiated as described earlier. The refolding reactions were initiated in buffer A that were preincubated at different temperatures (15°C, 20°C, 25°C, 30°C and 35°C) and monitored by measuring the recovery of GFP fluorescence. Rates obtained by fitting the refolding traces obtained at different temperatures are plotted against (1/T) for sGFP (grey circles), and Wt GFP (black circles). The rates were obtained from independent replicates of 3 refolding reactions at each temperature. Line plots are the fits obtained by fitting the temperature-dependent refolding rates to the Arrhenius equation as described in the supplemental text. (E) The calculated enthalpy of activation (ΔH#), and ρ obtained from Arrhenius fitting of sGFP and Wt GFP refolding shown in (D). (F) Close up of K45 (orange:side-chain carbons of K45, navy blue:εN) region on GFP (using 1GFL) (Yang et al., 1996), and its interacting amino acids. The model was prepared using Chimera. (G) Comparison of the refolding rate of sGFP in the presence of different salts of equal ionic strength. Spontaneous refolding of sGFP was initiated by a 100-fold dilution of the unfolded proteins in buffer A at 25°C. For salt assisted refolding unfolded protein was diluted 100-fold in buffer A containing different salts (100 mM MgCl_2_, or 300 mM NaCl, or 300 mM KCl) of equal ionic strength so that the final concentration of unfolded protein is 200 nM. Recovery of GFP fluorescence over time was followed to monitor refolding. See Figure S2 and S3.

To understand the type of folding-barrier incorporated by the K45E mutation in sGFP, we analyzed the Arrhenius plots for the folding of sGFP and Wt GFP (Figure 2D, Table S1). We fitted temperature-dependent refolding rates of the proteins using a full model of the Arrhenius equation containing the ΔCp^#^ term (to account for the difference in heat capacity of the folding intermediate and the transition state) (Dandage et al., 2015). sGFP did not have a temperature-dependent slope suggesting the absence of any enthalpic folding barrier (ΔH^#^) while Wt GFP had a high ΔH^#^ indicating an enthalpic barrier to folding (Figure 2E, Table S1). Since sGFP refolded slower than Wt GFP, this suggested that the rate-limiting barrier to folding in sGFP is non-enthalpic (Figure S3A). Complementary to ΔH^#^ is the value of ρ that is a composite function of the frequency across the transition barrier (prefactor for Arrhenius function) and the entropic assistance towards reaching the transition state (TS) (Figure S3A); lower ρ is associated with higher entropic barrier (Dandage et al., 2015). ρ was higher for Wt GFP compared to sGFP (Figure 2E, Table S1) demonstrating that sGFP faced an entropic barrier in folding trajectory. This was consistent with earlier findings with other GroEL substrates (Chakraborty et al., 2010; Georgescauld et al., 2014). sGFP was likely trapped in a folding intermediate I_1_ that was stabilized by flexible regions that prevented the formation of native contacts. Analysis of GFP structure revealed that the residue mutated in sGFP, K45, interacted with multiple negatively charged residues in Wt GFP (E32, D210, and E213) (Figure 2F). This lysine residue also formed a core for nucleating long-range interactions in a loop region. The substitution of lysine (K45) to a negatively charged glutamate in sGFP could have led to the loss of these interactions and introduced repulsive interactions that would destabilize this region. Spontaneous refolding rate of sGFP increased with an increase in salt concentration (salt shields charges and prevents repulsive and attractive interactions) (Figure 2G) and charge repulsion in sGFP likely prevented folding by increasing flexibility around this region. Similar salt-dependence was absent in Wt GFP (Figure S2E) demonstrating that substitution of lysine with glutamate introduced electrostatic repulsions that limit sGFP refolding. Taken together, the folding landscape of sGFP had a rate-limiting entropic trap driven by flexibility of I_1_ that accounts for the slower folding rate of sGFP compared to Wt GFP.

### GroEL/ES alters the folding pathway of sGFP

To check if the folding of sGFP was altered by GroEL/ES system, we tested for its chaperone-dependence *in vitro* by reconstituting chaperone-assisted refolding reactions. The refolding of sGFP was strongly accelerated (3∼4 fold) by GroEL/ES (Figure 3A) system while that of Wt GFP remained unchanged (Figure 1A, lower panel) at 25°C. Of importance, the *in vivo* chaperonin-dependence of sGFP was found to be intact even at 25°C, the temperature used for *in vitro* chaperone-assisted refolding experiments (Figure S3B). Chaperonin-dependent acceleration of sGFP refolding was reliant on the presence of all the components; GroEL, GroES, and ATP (Figure 3B). Thus, like authentic class-III (or IV) substrates (Kerner et al., 2005; Niwa et al., 2016) this lab-evolved *in vivo* substrate depended on the full-cycle of the GroEL/ES system. Moreover, the DnaK/J/E system did not alter the refolding rate of sGFP *in vitro* (Figure S3C) in line with the *in vivo* data described earlier (Figure 1G). This demonstrated that the isolated mutant sGFP, was a specific substrate of GroEL/ES *in vivo* and *in vitro*, and GroEL/ES accelerates the refolding rate of sGFP. A single mutation K45E on Wt GFP conferred GroEL/ES dependence for its *in vivo* folding as well as accelerated refolding rate *in vitro*.

**Figure 3:**
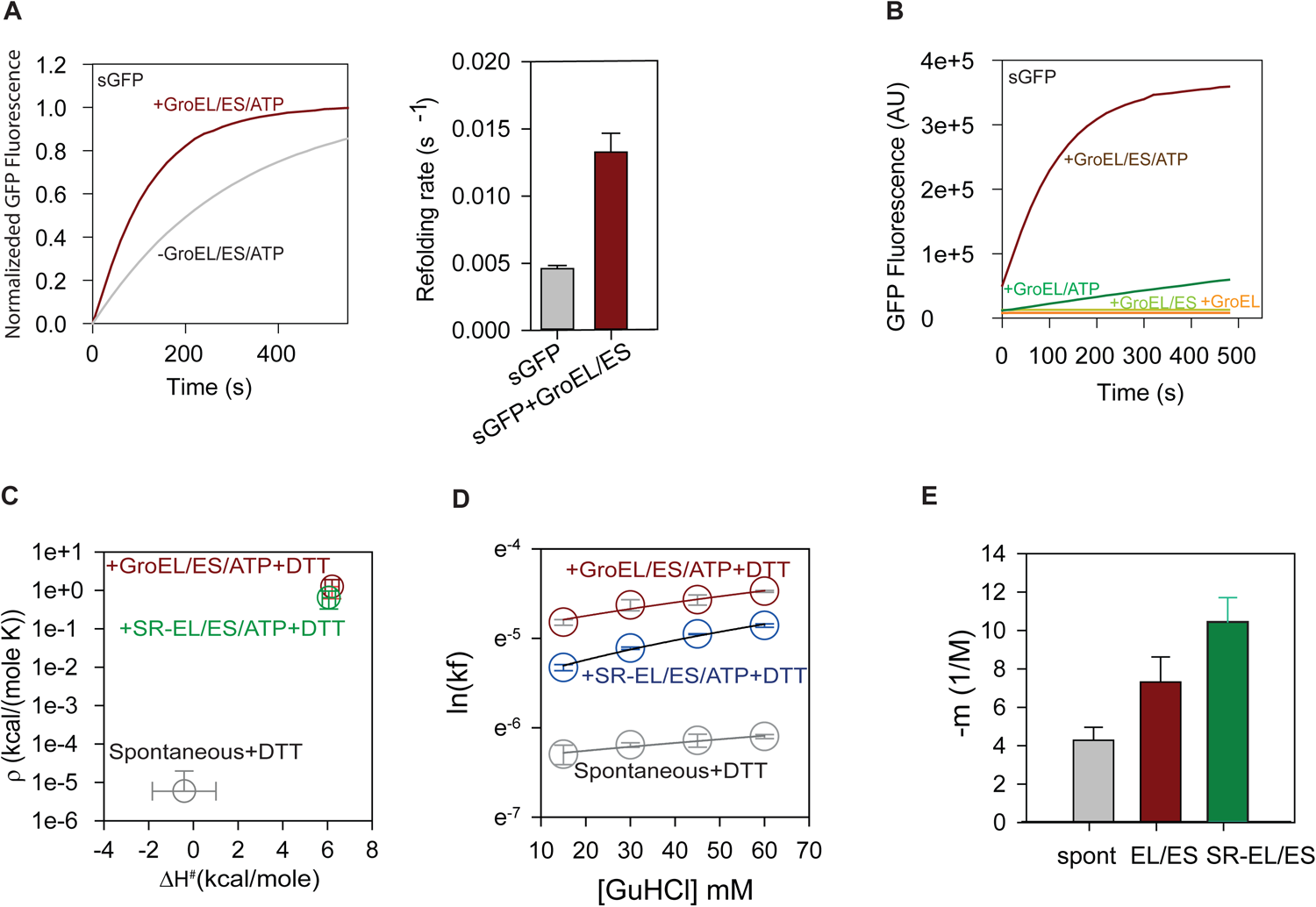
GroEL/ES alters the folding pathway of sGFP. (A) Left panel: Comparison of the refolding kinetics of sGFP in the presence and absence of GroEL/ES/ATP. Spontaneous refolding of sGFP was initiated as mentioned earlier either in buffer A. For GroEL/ES assisted refolding unfolded protein was diluted 100-fold in buffer A containing 400 nM of GroEL (all concentrations of GroEL are in terms of tetradecamer), 800 nM GroES (all the concentrations of GroES are in terms of heptamer) so that the final concentration of unfolded protein is 200 nM. Subsequently, refolding was started by adding 2 mM ATP. Recovery of GFP fluorescence over time was followed to monitor refolding. Right panel: Refolding traces were fitted to single exponential kinetics to obtain the refolding rates shown as bar plots. Three independent refolding runs were fitted separately to obtain the standard deviation in rates (shown as errors on the bar graph). (B) Monitoring the dependence of sGFP refolding on the different components of EL/ES. Unfolded sGFP was diluted 100-fold either in the presence of GroEL (400 nM), or GroEL (400 nM) + GroES (800 nM) or GroEL (400 nM) + ATP (2 mM) or GroEL (400 nM) + GroES (800 nM) + ATP (2 mM) and refolding was monitored by the recovery of GFP fluorescence. (C) The calculated enthalpy of activation (ΔH^#^), and ρ obtained from Arrhenius fitting of GroEL/ES/ATP and SR-EL/ES/ATP assisted refolding of sGFP. (D) Effect of GuHCl on refolding rate upon spontaneous, GroEL/ES/ATP and SREL/ES/ATP assisted refolding of sGFP. The rates [ln(k_f_)] of spontaneous, GroEL/ES/ATP, and SREL/ES/ATP assisted refolding of sGFP at different GuHCl concentration are plotted against different GuHCl concentrations. (E) The m-value (dependence of refolding rate on GuHCl concentration) of folding upon spontaneous, GroEL/ES/ATP, and SR-EL/ES/ATP assisted refolding of sGFP.

To find the mode of acceleration by GroEL/ES, we obtained the ΔH^#^ and ρ for GroEL/ES assisted folding from modified Arrhenius analysis (Dandage et al., 2015). Notably, the GroEL/ES system did not decrease the enthalpic barrier (ΔH^#^) thereby working through the route of entropic destabilization (Figure 3C, Table S1). A higher value of ρ for GroEL/ES assisted refolding than for spontaneous refolding indicated a lower entropic barrier to refolding in the presence of GroEL/ES (Figure 3C, Table S1). This was consistent with other substrates of the GroEL/ES system (Chakraborty et al., 2010; Georgescauld et al., 2014) suggesting that entropic traps in folding landscapes may characterize a GroEL/ES substrate in general, and GroEL/ES could have a general role of affecting entropic destabilization to assist protein folding. The difference in GroEL/ES-assisted and the spontaneous refolding pathway further corroborated with the difference in m-value (dependence of refolding rate on GuHCl concentration) of folding (Figure 3D and 3E).

To check if the allosteric cycle of GroEL/ES or iterative annealing played a role in accelerating sGFP refolding, we used the single-ring version of GroEL (SR-EL) (Horwich et al., 1998) that lacks the negative allostery between rings, encapsulates the substrate and completes folding inside the cavity. SR-EL/ES could accelerate the refolding of sGFP like GroEL/ES *in vitro* (Figure S3D) and the Arrhenius parameters (Figure 3C) and m-value (Figure 3D and 3E) from SR-EL/ES assisted refolding were similar to that assisted by GroEL/ES. The similarity of Arrhenius parameters between SR-EL/ES and GroEL/ES dependent folding of sGFP suggested that the temperature dependence of the refolding rates were independent of the temperature effects on GroEL/ES allostery and was primarily determined by the folding landscape of sGFP. The similarity of the Arrhenius parameters and the m-value suggested that the folding mechanism of sGFP had maximal contribution from the encapsulated folding environment in the cage that had the potential to change the folding landscape. Thus, a decrease in the entropic barrier of the refolding-substrate was primarily affected by the GroEL/ES cavity and could rescue the entropically trapped state of the mutant GFP.

### sGFP refolding is also limited by a non-native contact formation

Interestingly the yield of spontaneous refolding of sGFP drastically reduced in the absence of any reducing agent in the refolding buffer (Figure 4A) and reduced further in the presence CuCl_2_, a disulfide-assisting catalyst (Figure S4A). Since each molecule of sGFP has two cysteines (C48 and C70) that are distal in the native structure (Figure S4B), there were two possibilities, 1) the proteins formed intermolecular-disulfides and formed covalently bonded aggregates or 2) the proteins formed an intramolecular-disulfide bond that trapped the molecules in the non-native state. Formation of an intramolecular-disulfide would result in the co-migration of two peptides that were disulfide-bonded if the protein was digested with trypsin (Figure 4B). Mass spectrometry confirmed the presence of the expected product during sGFP refolding in non-reducing conditions and proved the formation of an intramolecular-disulfide (Figure 4C and S4C); this peptide was not detectable in the presence of a reducing agent. Consistently, we observed the non-disulfide bonded peptide fragments in the presence of DTT (Figure 4D). This peptide was ∼100 fold enriched in the presence of DTT than in its absence (Table S2). This proved that the intermediate was trapped with an intramolecular-disulfide while refolding spontaneously in non-reducing conditions. Non-reducing gel electrophoresis confirmed that intermolecular-disulfide was not a major species when refolding was initiated in the absence of DTT (Figure 4E). Indicating that an intramolecular-disulfide trapped the folding intermediate in a non-native state and not an intermolecular-disulfide. Thus, the most likely model for spontaneous refolding of sGFP was through a refolding intermediate I_1_ that was free to fold to its native state with an apparent rate of ∼2e-3 s^-1^. In oxidizing conditions, I_1_ converted to a quasi-stable species with non-native contacts (I_1_*) which rapidly converted to the terminally misfolded intramolecular-disulfide bonded species (I_2_) (Figure 4F, and S4D).

**Figure 4:**
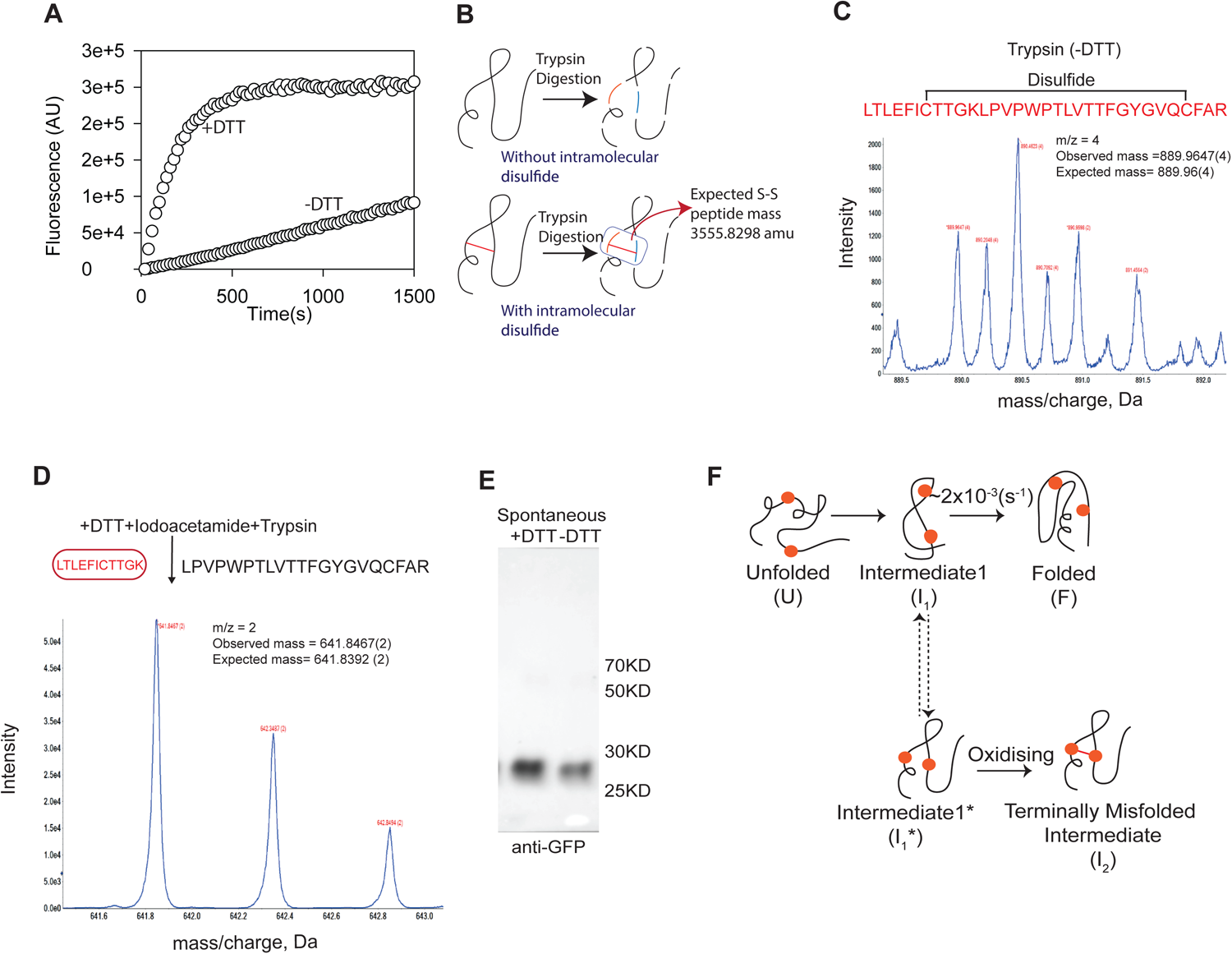
GroEL/ES prevents non-native contact formation. (A) Refolding of sGFP was initiated in buffer A (shown as +DTT) or buffer B (20 mM Tris, 20 mM KCl, 5 mM MgCl_2_, pH 7.4) (shown as -DTT). Refolding was monitored by the recovery of GFP fluorescence over time. (B) Schematic representation of a hypothetical fragment pattern when an intramolecular disulfide is formed (in the absence of DTT) and in its absence (in the presence of DTT). (C) The expected peptide sequence and mass upon the formation of the intramolecular disulfide is shown along with the experimentally observed isotopic mass distribution obtained for the peptide when refolding of sGFP is initiated in the absence of DTT. (D) The peptide fragment expected when disulfide bonding is prevented during sGFP refolding in DTT, and the experimentally observed isotopic mass distribution for the peptide is shown. (E) Non-reducing PAGE to monitor the formation of intermolecular disulfide bonds. Folding of sGFP was initiated as described earlier in buffer B or buffer A at a final concentration of 500 nM sGFP. The proteins were then boiled and loaded onto a non-reducing polyacrylamide gel to resolve the proteins. Immunoblotting was performed to detect sGFP using anti-GFP antibody. (F) A simplistic model of the spontaneously refolding sGFP. The unfolded state reaches the folding intermediate I_1_ which then converts to the native state N. I_1_ is in rapid equilibrium with I_1_* where the cysteines (shown as orange circles) come close. I_1_*, in turn, can form the disulfide in oxidizing conditions and form a terminally misfolded state I_2_ that is refractile to refolding even in the presence of DTT. See Figure S4.

Since sGFP folds poorly in the absence of DTT, we asked if delayed addition of DTT could rescue the disulfide-trapped refolding intermediate I_2_. When the refolding was initiated in the presence of 2 µM CuCl_2_, the delayed addition of DTT could not restore the sGFP folding (Figure 5A), suggesting that sGFP in the disulfide-bonded state enters an irreversibly misfolded state I_2_ that was refractile to reducing agents. Since more than 95% of the molecules reached I_2_ within 5 minutes of the start of the refolding reaction, it revealed that there was a rapid equilibrium between I_1_ and I_1_* (Figure 4F, dotted arrows) that was much faster than the time scale of folding. Suggesting that the formation of I_1_* did not affect the apparent rate of folding from I_1_ to N. Additionally, under conditions that favor disulfide formation, sGFP folding was limited by an enthalpic trap that was driven by non-native contact formation in the regions surrounding the two cysteines. The traps for this substrate could be switched on/off: the enthalpic trap could be switched off by DTT (reducing agent), and the entropic trap could be attenuated in the presence of salts.

**Figure 5:**
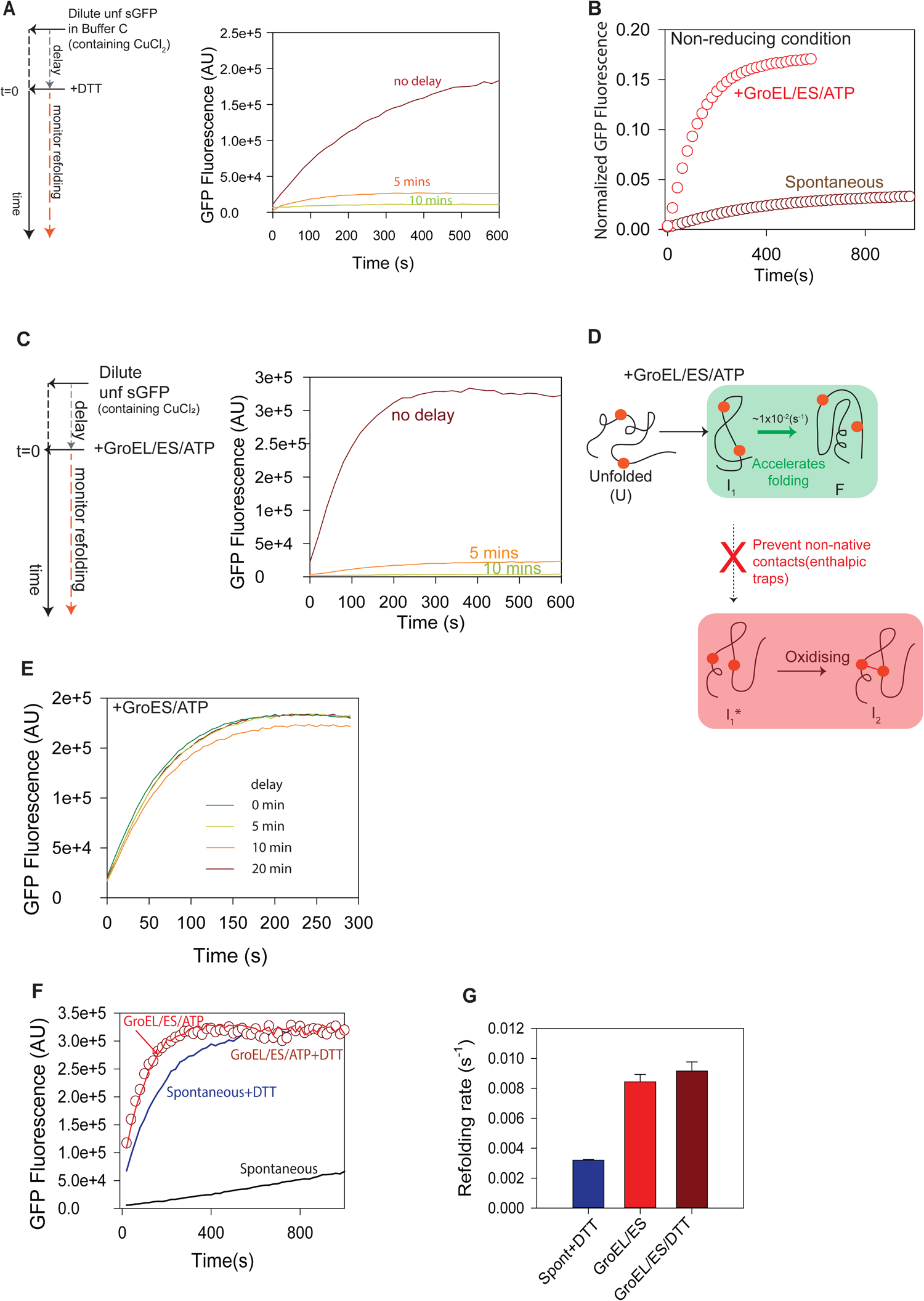
GroEL/ES accelerates refolding rate by decreasing the entropic barrier. (A) Effect of delayed addition of DTT to spontaneously refolding sGFP. sGFP refolding was initiated in buffer C (20 mM Tris, 20 mM KCl, 5 mM MgCl_2_, 2 μM CuCl_2_ pH 7.4) at a final concentration of 200 nM of sGFP. DTT was either added before unfolded sGFP was diluted in the buffer to start refolding (no delay) or after 5 min or 10 min. Refolding traces for 5 min and 10 minutes delayed addition of DTT are shown from the time DTT was added. (B) Spontaneous or GroEL/ES dependent refolding of sGFP was initiated as described earlier, except that the refolding was performed in buffer B for both the refolding reactions. (C) Effect of delayed addition of GroEL/ES/ATP to spontaneously refolding sGFP. Refolding of sGFP was initiated as earlier in buffer C, and GroEL/ES/ATP (400 nM/800 nM/2 mM respectively) was added after a time-delay of 5 min or 10 min. For zero delay, GroEL/ES/ATP was added to buffer C before adding unfolded sGFP. (D) Schematic of the refolding pathway of sGFP in the presence and absence of GroEL/ES. GroEL/ES/ATP efficiently prevents the formation of I_1_* (shown by the red arrow) and also allows more efficient conversion of I_1_ to N (shown by the green arrow). (E) GroEL-binding is sufficient to prevent sGFP misfolding. sGFP refolding was initiated as earlier, in buffer C containing 400 nM GroEL. GroES/ATP (800 nM/2 mM, respectively) was added after time delays of 5 min, 10 min, or 20 min. Refolding was monitored although the time course, traces shown are after the addition of GroES/ATP. For zero delay addition, GroES/ATP was present along with GroEL in buffer C before unfolded sGFP was added to the buffer. (F) Effect of DTT on GroEL/ES assisted refolding rate of sGFP. Refolding of sGFP was initiated as described in buffer C (black line), buffer C containing GroEL/ES/ATP (400 nM, 800 nM, 2 mM, respectively) (red line), buffer A (blue line), and buffer A containing GroEL/ES/ATP (400 nM, 800 nM, 2 mM, respectively) (maroon circles). (G) Rates for refolding in the latter three (from Figure 5F) were obtained by fitting to the exponential rate equation and is shown as bar graphs. Errors shown are standard deviations over three independent replicates. See Figure S5.

### GroEL/ES accelerates refolding rate by decreasing the entropic barrier

Could the enthalpic traps generated by the formation of non-native contacts be rescued by GroEL/ES? Notably, the GroEL/ES system was able to accelerate the folding of sGFP and increase the yield of refolding even in the absence of DTT (Figure 5B). Since the cysteines were not proximal in the native structure, disulfide formation was dependent on non-native contact formation and hence misfolding. This suggested that GroEL/ES prevents the formation of a non-native contact during the conversion from I_1_ to I_2_. To further understand this, we performed the refolding with delayed addition of the GroEL/ES system after initiating the refolding of sGFP in the presence of 2 µM CuCl_2_ (Figure 5C). Delayed addition of GroEL/ES led to a near-complete loss of refolding, indicating that the GroEL/ES system was unable to break the preformed disulfide, as expected, and could only prevent the formation of the non-native disulfide by altering the folding path (Figure 5D). Taken together, the refolding intermediate (I_1_) formed non-native contacts to form a quasi-stable species (I_1_*) that rapidly formed non-native disulfide (under oxidizing conditions) to form the intermediate I_2_. This intermediate was terminally misfolded and refractile to folding assistance by DTT or GroEL/ES system. Implying that GroEL/ES system was able to prevent the formation of non-native contacts in I_1_ (Figure 5D, pink box) and hence the formation of I_2,_ channeling folding of sGFP through a productive route.

GroEL-binding is known to cause partial unfolding of proteins, and can potentially remove non-native interactions (Mapa et al., 2012; Sharma et al., 2008). To test this, we initiated the refolding of sGFP in the presence of GroEL in a non-reducing buffer, but in the absence of GroES or ATP. GroEL alone did not assist refolding (Figure S5A). Delayed addition of GroES/ATP restarted refolding with the same rate as observed when GroEL/ES/ATP is added without any delay (Figure S5B). In contrast to delayed addition of the full GroEL/ES/ATP system, the presence of GroEL prevented misfolding and the amplitude of refolding did not drop even after a 20 minute of delay in addition of GroES/ATP to GroEL-bound unfolded sGFP. (Figure 5E, and S5C). This proved that GroEL-binding to the non-native state of sGFP prevented the protein from forming non-native interactions, thereby maintaining it in a folding-competent state for a long time. Taken together, the GroEL/ES system or GroEL alone can prevent cysteines from coming in close contact; thereby averting the formation of non-native contacts that would result in enthalpic traps during folding. While GroEL/ES was able to prevent I_1_ to I_2_ conversion, the refolding rate of GroEL/ES assisted folding was same in the presence or absence of DTT in the refolding reaction (Figure 5F). Essentially, GroEL/ES assisted refolding rate was faster than the spontaneous folding rate of sGFP even in the presence of DTT (Figure 5G). The refolding rate of sGFP did not increase as a function of DTT concentrations in the range used for the refolding here (Figure S5D) negating the argument that partial reduction of disulfide was the reason for slower spontaneous folding than with GroEL/ES system. This clearly demonstrated that the GroEL/ES system could accelerate the refolding rate of sGFP even under (reducing) conditions where I_1_ to I_2_ conversion was completely prevented. Thus, in addition to preventing I_1_ to I_2_ conversion, GroEL/ES assistance increased the folding rate from I_1_ to N irrespective of the redox condition of the refolding reaction (Figure 5D). However, the enthalpic trap was not specific to the K45E mutation of GFP, rather it was the property of the GFP backbone that showed a redox-dependent change in refolding amplitude and rate (Figure S5E). Thus, the entropic component of the folding barrier possibly rendered the K45E mutation amenable to GroEL/ES dependent folding *in vivo* and *in vitro*.

Collectively, GroEL/ES worked in a bipartite manner to assist sGFP refolding by (Figure S3A, upper panel), 1) preventing non-native contact formation and hence the formation of non-productive off-pathway intermediates that had the potential to form a terminally misfolded conformation and 2) decreasing the entropic barrier in the folding pathway to increase the folding rate (Figure S3A, lower panel).

### Implications

While proteins evolve, they accumulate mutations. A subset of these mutations may enhance the existing activity or impart new functions, they are also more likely to destabilize proteins than mutations on other regions of the protein surface (Tokuriki et al., 2008). Chaperones have been proposed to aid these transitional sequences allowing them to cross fitness barriers.

Aiming to link the molecular mechanism of chaperone-dependent buffering to the specific perturbations in protein-folding landscapes we used GroEL/ES as the model chaperone. We found that the GroEL/ES system buffers the entropic traps that can arise due to mutations that stabilize the folding intermediates entropically.

Interestingly, GroEL/ES could also prevent the formation of incorrect contacts in the folding polypeptide (rescue of enthalpic traps). However, this was not an exclusive activity of GroEL/ES as we show that even the DnaK/DnaJ/GrpE chaperone machinery shares this capability. Given that binding of non-native proteins with DnaK (Banerjee et al., 2016; Mattoo et al., 2014; Rüdiger et al., 1997) or DnaJ (Tiwari et al.) or GroEL (Lin et al., 2008; Sharma et al., 2008) can unfold protein chains and melt preformed contacts, it is conceivable that the holdase and the unfoldase action is required for removing enthalpic traps from folding landscapes. DnaK machinery in *E. coli* cytosol is more abundant than the GroEL/ES machinery and hence the mutations that introduce enthalpic traps in the folding pathway are more likely to be channeled through the former for efficient rescue.

While the current knowledge can be used in protein redesigning, our proposition, that GroEL/ES may specifically buffer mutations that traps flexible folding intermediates in entropic traps, will have interesting implications if it is found to be generally true in natural evolution.

## METHODS

### EXPERIMENTAL MODEL AND SUBJECT DETAILS

#### Strains, Plasmids, and Proteins

*E. coli* strain DH5α was used for cloning, WT *E. coli* K-12 (BW25113) strain was used for expression of arabinose inducible pBAD GFP and BL21 (DE3) was used for protein expression and purification. Protein concentrations were determined spectrophotometrically at 562 nm using BCA kit (Pierce-ThermoFisher Scientific). Deletion strains were obtained from CGSC as part of Keio collection (Baba et al., 2006).

### METHOD DETAILS

#### Construction of mutant GFP library

Mutant GFP library was made in arabinose inducible pBAD vector using a random mutagenesis approach by error-prone polymerase Mutazyme II (Agilent) that incorporated 7 to 11 mutations per kb of the template. The said library has a total complexity of around 10,000 mutants. The reporter is constructed such that GFP and mCherry are under the same arabinose inducible pBAD promoter in an operon to give a readout of GFP according to the mutation created on it but the mCherry readout will remain similar thus serving as an internal control for transcription, translation, and inducibility.

#### Screening of mutant GFP library for GroEL/ES dependent GFP mutant

Wild type *E. coli* cells (WT) were transformed with the mutant GFP library maintaining 10-fold converge for preserving complexity. 0.4 OD600 cells were induced with 0.1% arabinose and fluorescence was observed three hours post-induction at 37°C after diluting cells in 1X PBS and incubating at 37°C for 1 hour. Fluorescence of the mutant library was studied in a pooled manner against wild type GFP. The entire library was sorted into populations of very low fluorescent, low fluorescent, and mildly less fluorescent according to the GFP fluorescence. Each of these populations was purified and plasmids prepared. WT cells were co-transformed with GroEL/ES over-expressing plasmid (Castanie et al., 1997) and pool of GFP mutant plasmids in a sequential manner, maintaining the minimum 10X coverage. The pool of transformants were grown and GroEL/ES induced by adding 0.5 mM IPTG 30 minutes before induction of GFP by 0.1% arabinose. After that the induced cultures were grown for another 3 and half hours. FACS was performed to sort *E*.*coli* cells showing high fluorescence upon GroEL/ES overexpression. To confirm the dependence of sorted clones on GroEL/ES, plasmid pool was prepared from the sorted cells (harboring pBAD GFP as well as pOFX GroEL/ES) and digested using SacII (linearizes pOFX GroEL/ES only) to obtain GFP clones post-transformation in wild type *E*.*coli* cells. Single clones of GFP were picked from here and checked for their fluorescence in the presence and absence of GroEL/ES overexpression. The isolated mutant having higher fluorescence in GroEL/ES overexpression was identified by Sanger sequencing. Isolated GroEL/ES dependent mutant of GFP (K45E) was cloned in pET SUMO between BamHI and HindIII restriction sites and purified using *E. coli* BL21 (DE3) for further characterization.

#### *E.coli* GroEL/ES, DnaK, DnaJ, GrpE Expression, and Purification

GroEL/ES, GroEL chimeras, DnaK, DnaJ, GrpE were purified using *E. coli* BL21 (DE3) as described (Kerner et al., 2005; Mapa et al., 2012; Tiwari et al., 2013). GroEL/ES was expressed from pOFX plasmid for co-expression studies (Castanie et al., 1997).

#### Solubility of sGFP *in vivo*

WT *E.coli* K-12 (BW25113) cells containing pBAD sGFP were transformed with pOFX GroEL/ES. 0.1% inoculation was done in 10 ml LB medium added with chloramphenicol (35 μg/ml) and ampicillin (100 μg/ml) from overnight grown cultures and grown till OD600-0.5 at 37°C, 200 rpm. 0.5 mM IPTG was added to cells with pOFX GroEL/ES 30 minutes before the induction of sGFP with 0.1% arabinose for 3 hours. The cell type in which we transformed only pBAD sGFP were directly induced with 0.1% arabinose for 3 hours after reaching OD600-0.5. Cells were harvested at 4000 rpm for 10 minutes and resuspended in 1 ml PBS pH 7.4, 2 mM DTT. Lysis was done using sonication followed by high-speed centrifugation to separate soluble and pellet fraction which was separately loaded onto 12% SDS-PAGE. Gel visualized by Coomassie staining.

#### Spontaneous and chaperonin assisted *in vitro* refolding of sGFP

Wt GFP and sGFP (20 μM each) were denatured in buffer containing 6 M GuHCl in buffer A (20 mM Tris, 20 mM KCl, 5 mM MgCl_2_, 2 mM DTT, pH 7.4) for 1 hour at 25°C and refolded upon 100 fold dilution into buffer A. Either of three refolding buffers were used for refolding, buffer A to mimic reducing conditions, buffer B (20 mM Tris, 20 mM KCl, 5 mM MgCl_2_, pH 7.4) to mimic non-reducing conditions and buffer C (20 mM Tris, 20 mM KCl, 5 mM MgCl_2_, 2 µM CuCl_2_, pH 7.4) to mimic oxidizing conditions.

GroEL/ES assisted refolding was done in the presence of (400 nM) GroEL (tetradecamer), (800 nM) GroES (heptamer), (substrate:GroEL:ES :: 1:2:4) and the refolding was started by addition of 2 mM ATP. SR-EL/ES assisted refolding was done in the presence of (800 nM) of SR-EL (heptamer) (800 nM) GroES (heptamer) and the refolding was started by the addition of 2 mM ATP. GFP fluorescence at 480 nm excitation (slit width 2 nm) and 515 nm emission (10 nm slit width) was monitored as a readout of refolding using Fluorolog 3 Spectrofluorometer (Horiba). Buffer conditions are described in the figures. All the unfolding and refolding experiments were carried out at 25°C unless specified.

#### GuHCl concentration-dependent spontaneous and chaperonin assisted *in vitro* refolding of sGFP

sGFP (80 μM) were denatured in buffer containing 6 M GuHCl in buffer A for 1 hour at 25°C and was refolded upon 400 times dilution in buffer A alone or buffer A containing (400 nM) GroEL (tetradecamer), (800 nM) GroES (heptamer) and the refolding was started by addition of 2 mM ATP. GuHCl present in unfolded sGFP was diluted to 15 mM upon 400 times dilution in buffer A. GuHCl concentration was increased to 30 mM, 45 mM, 60 mM by adding GuHCl from outside in refolding mixture.

#### Analysis of temperature-dependent refolding using Arrhenius equation

To obtain thermodynamic parameters that define the barrier between the refolding intermediate I_1_ and the transition state (TS) of folding we used the following equation essentially as defined in (Dandage et al., 2015).

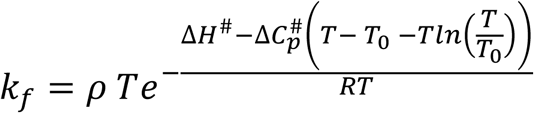

Where

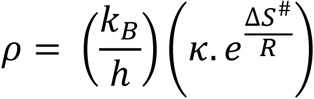

and k_f_ is refolding rate, R is the universal gas constant, ΔH^#^, ΔS^#^ and ΔC_p_^#^ are the differences in enthalpy, entropy and heat capacity at constant pressure and the reference temperature T_0_ between TS and I_1_, respectively, T is the temperature of refolding reaction, T_0_ is the reference temperature at which the parameters are determined (here itis 298.15 K). κ is the transmission factor, that reports the proportion of activations that lead to the formation of the native state (N), and k_B_ and h are Boltzmann and Planck constant respectively. Since ΔS^#^ and A linearly combine, it is not possible to obtain independent estimates of these two parameters by non-linear regression. Hence, we combine it to obtain ρ. This term indicates the ease of barrier crossing, either because the diffusion is faster or because the barrier is less broad (due to lower ΔS^#^). A high ΔH^#^ indicates a higher enthalpic barrier, and a low ρ indicates an entropic (diffusion-limited) barrier. The equations were fitted using standard non-linear regression fitting using R or Octave. Fitting was performed by varying the starting parameters by 4-fold within the rage of expected values reported for globular proteins (Dandage et al., 2015), and the fitting was deemed satisfactory only when the r-squared values were above 0.9 and the dependencies were above 0.9 for the different parameters that were floated during fitting. The floating parameters were, ρ, ΔC_p_, ΔH.

#### Mass spectrometry analysis for disulfide bond detection

5 µg of control or DTT treated sGFP was digested by using sequencing grade trypsin (1:10 ratio, Trypsin:protein) for 16-18 hours at 37°C. Before the digestion DTT treated sample was alkylated by using 55 mM iodoacetamide. Tryptic digested peptides were reconstituted in 5µl of LC-MS grade water containing 0.1% formic acid and run on a quadrupole-TOF hybrid mass spectrometer (TripleTOF 6600, Sciex, USA) coupled to a nano-LC system (Eksigent NanoLC-400). Two microliters of sample was injected and loaded onto a reverse-phase peptide Chromo LC trap (200 mm 0.5 mm) column and peptides were separated using a C18 column (75 mm 15 cm, Eksigent). The samples were run using a gradient method using buffer X (99.9% LC-MS water + 0.1% formic acid) and buffer Y (99.9% acetonitrile + 0.1% formic acid). The gradient consists of 95% of buffer X for 2 minutes, and then shifted to 90% of buffer X for 8 minutes, and then decreased to 20% of buffer X in 42 minutes and finally shifted to 95% of buffer X again for 16 minutes at a consistent flow rate of 250 nl min-1. Data was acquired with a NanoSpray source installed in the TripleTOF 6600 System using a nebulizing gas of 20 psi, a curtain gas of 25 psi, an ion spray voltage of 2000 V, and a heater interface temperature of 75 0C. Information-dependent acquisition (IDA) mode was set up with a TOF/MS survey scan (350–1600 m/z) with an accumulation time of 250 ms. For fragmentation, a maximum of ten precursor ions per cycle was selected with each MS/MS spectrum (200–1800 m/z) accumulated for 70 ms with a total cycle time of approximately 2.05 seconds. Parent ions with a charge state from +2 to +5 and an abundance of more than 150 cps were selected for MS/MS fragmentation. Once an ion had been fragmented by MS/MS, its mass and isotopes were excluded for 3 seconds. The wiff files generated from Triple TOF 6600 (which contain both MS and MS/MS spectra) were analyzed using the protein pilot v5.0 for the identification of our desire protein. For disulfide bond detection and further spectra analysis was done using Biopharma view v2.0 and peak view 2.2 software (Sciex).

### QUANTIFICATION AND STATISTICAL ANALYSIS

Student t-test and R package for non-linear regression was used for statistical analysis. Flow-cytometry data was analyzed using octave.

### DATA AND SOFTWARE AVAILABILITY

All data are provided in the manuscript. We did not develop any new software.

### KEY RESOURCE TABLE

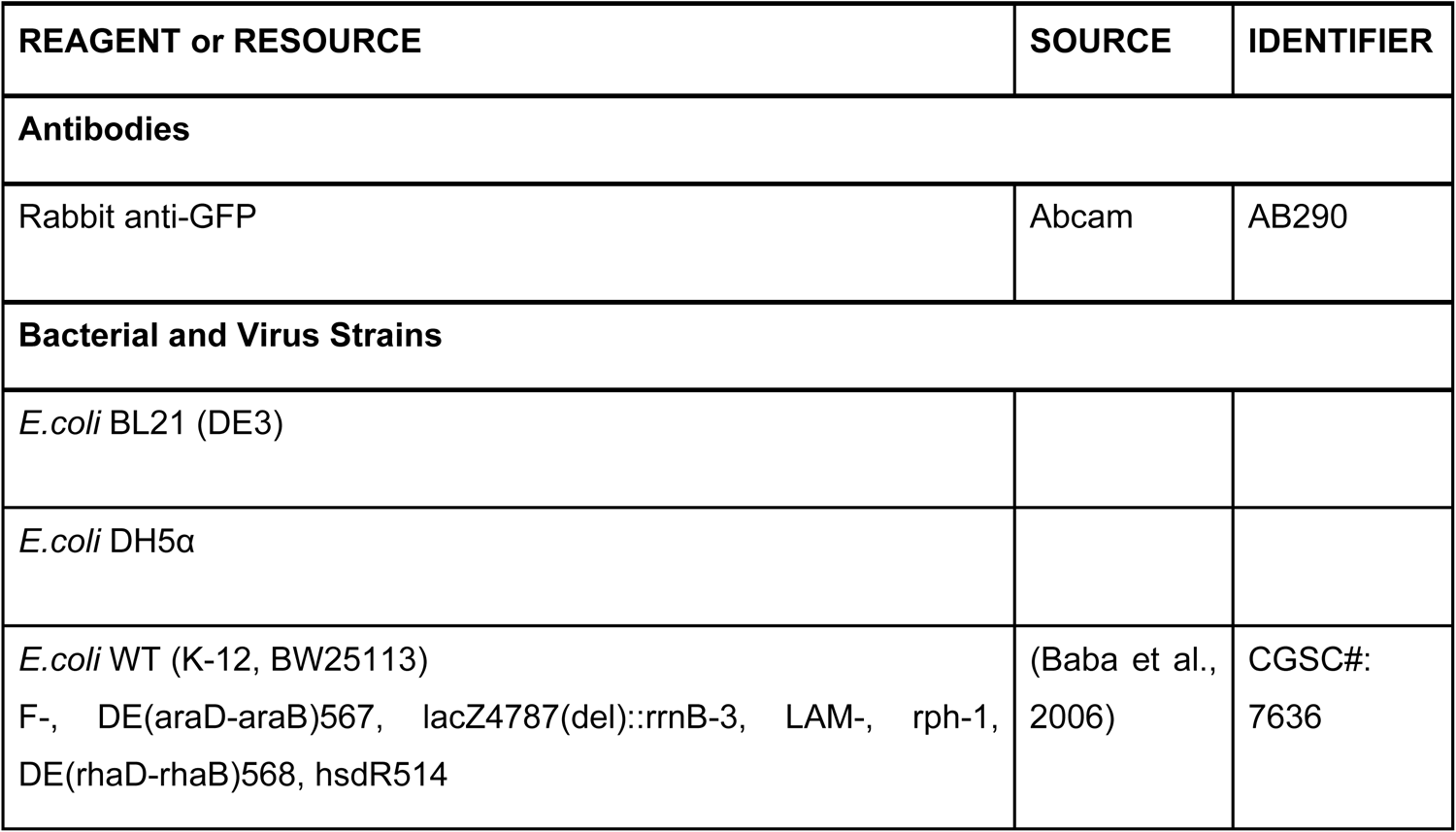
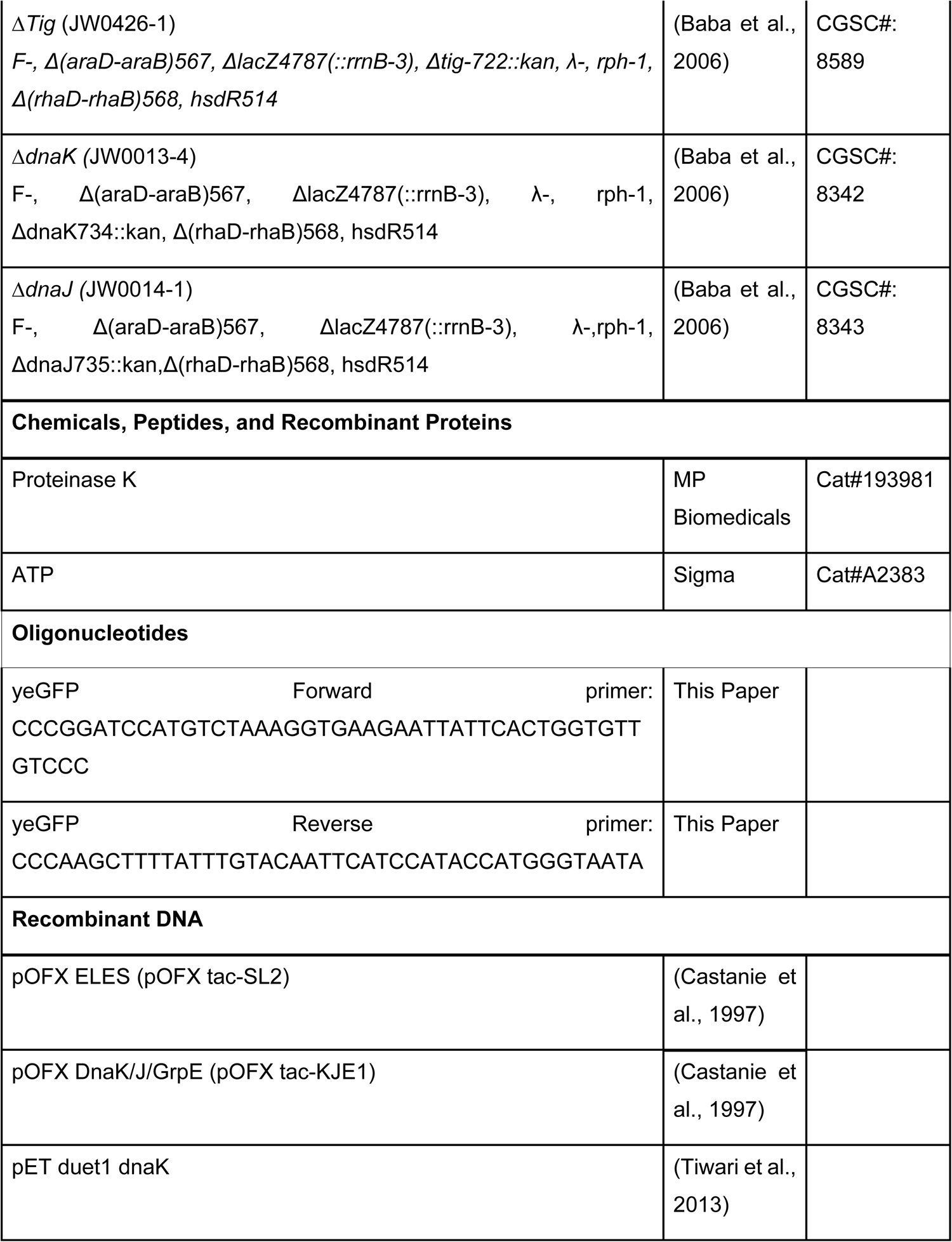
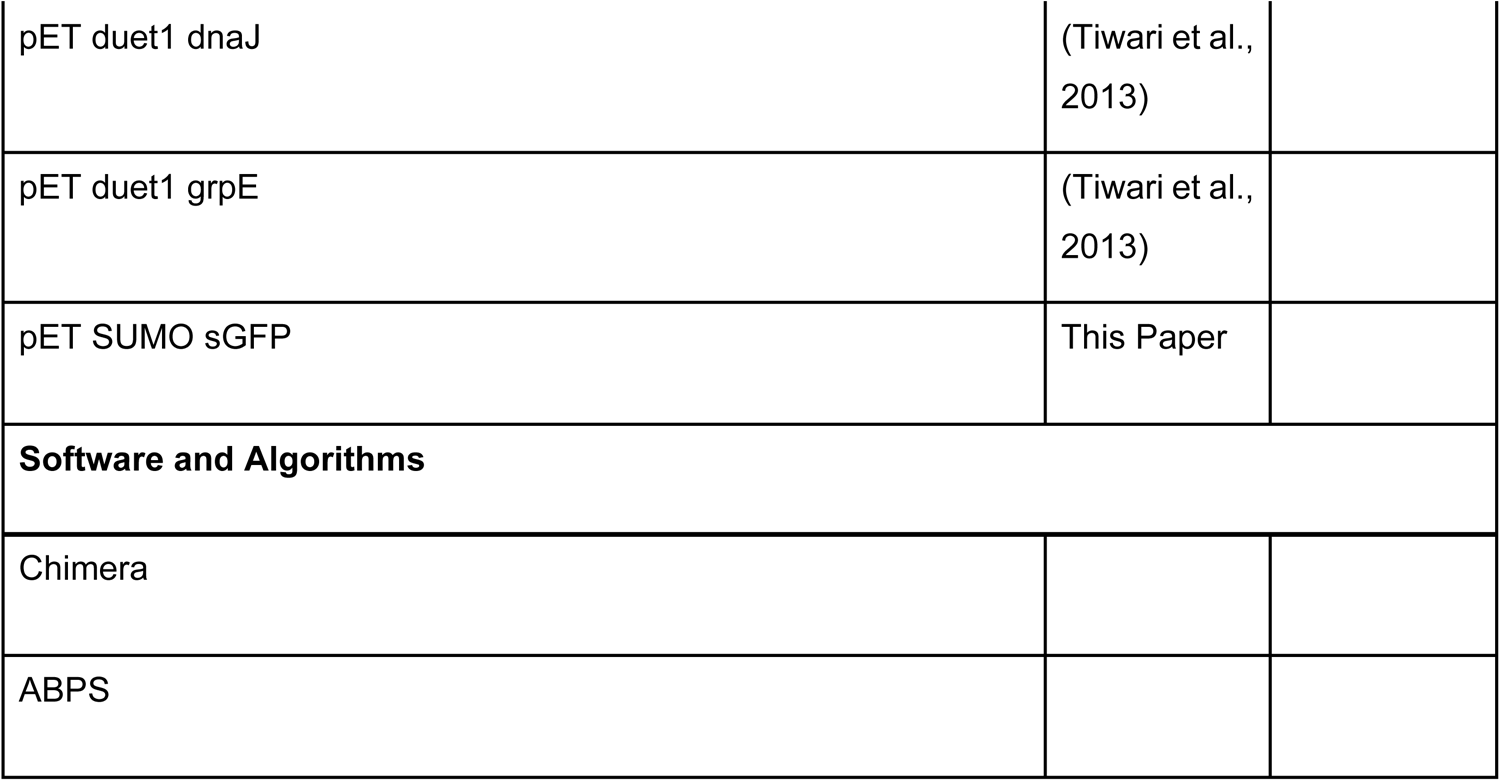

## ACKNOWLEDGEMENT

We thank Prof. Hideki Taguchi for the kind gift of pET21 SR1 plasmid. We also thank Kanika Saxena for sharing the starting plasmid with GFP and mCherry. KM acknowledges the funding from Department of Biotechnology (DBT), Government of India, grant number (BT/PR28386/BRB/10/1671/2018) and Science and Engineering Research Board (SERB), Government of India, for Core Research Grant (SERB/CRG/2019/006281) and SNU core funding. The work in KC Lab was supported by Swarnajayanthi Fellowship Grant from DST, and partly from BSC0124 from CSIR. Instrument support was also obtained from Wellcome Trust-DBT India Alliance and CSIR. KM and MA acknowledge SNU for infrastructural support and KC acknowledges CSIR and CSIR-IGIB for infrastructural support. A. Sadat and ST acknowledges DBT; AR, KV, AC, A. Singh acknowledge CSIR; AG and RC acknowledge UGC for fellowship support. MA acknowledges SNU doctoral fellowship.

## AUTHOR CONTRIBUTION

Conceptualization: Koyeli Mapa

Supervision: Koyeli Mapa, Kausik Chakraborty

Reagent Generation: Satyam Tiwari, Kanika Verma, Anwar Sadat, Aseem Chaphalkar,

Experiments: Anwar Sadat, Satyam Tiwari, Kanika Verma, Mudassar Ali, Vaibhav Upadhyay, Anupam Singh, Aseem Chaphalkar, Asmita Ghosh, Rahul Chakraborty, Kausik Chakraborty

Analysis: Arjun Ray (computational), Anwar Sadat (biophysics), Satyam Tiwari (biophysics), Kanika Verma (FACS), Rahul Chakraborty (MS), Kausik Chakraborty, Koyeli Mapa

Manuscript writing: Koyeli Mapa and Kausik Chakraborty with inputs from Anwar Sadat, Kanika Verma, and Aseem Chaphalkar. All the authors read and edited the manuscript.

## Supplementary Figures

**Figure S1:**
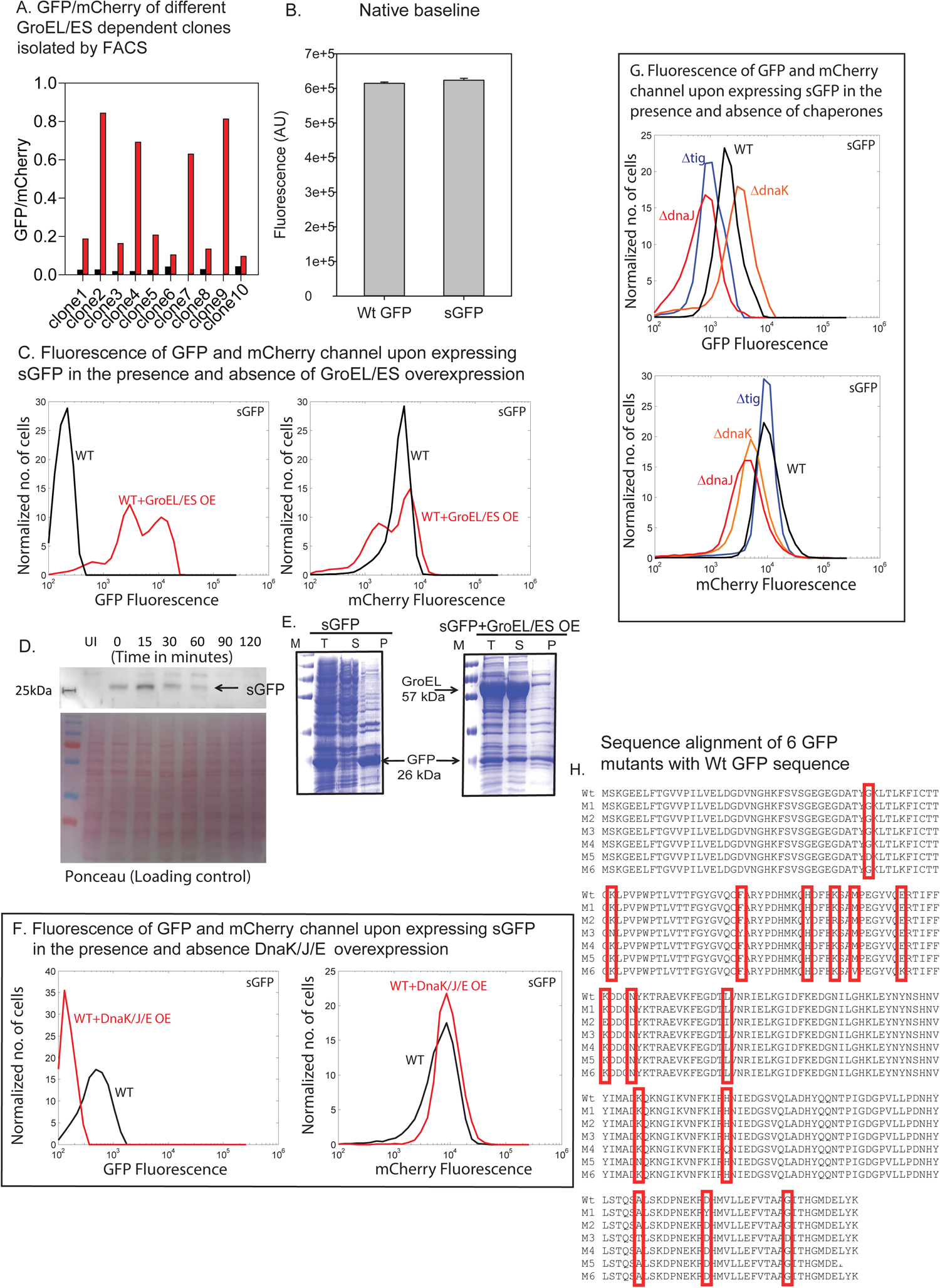
related to Figure 1. (A) Bar graph for *in vivo* fluorescence of GroEL/ES dependent 10 single GFP clones isolated through FACS with (red) and without (black) GroEL/ES overexpression. (B) Bar graph for native fluorescence of sGFP (K45E) compared to Wt GFP. (C) Independent fluorescence of GFP and mCherry channel for sGFP with (red) and without (black) GroEL/ES overexpression. GroEL/ES overexpression leads to an increase in GFP fluorescence (left panel) with similar mCherry fluorescence (right panel). (D) Western blot showing the degradation of sGFP after translation halt in the absence of GroEL/ES overexpression. Over time slow folding sGFP is degraded after translation is halted in absence of GroEL/ES overexpression. The western blot is representative of three independent experiments. (E) *in vivo* solubility of sGFP in the absence (left panel) and presence (right panel) of GroEL/ES over-expression. Partitioning of sGFP is shown in the total lysate (T), pellet fraction (P), and soluble fraction (S). (F) Independent fluorescence of GFP and mCherry channel for sGFP with (red) and without (black) plasmid-based DnaK/J/GrpE overexpression. Overexpression of DnaK/J/GrpE decreases GFP fluorescence (upper panel) with no effect on mCherry fluorescence (lower panel). (G) Independent fluorescence of GFP (upper panel) and mCherry (lower panel) channel for sGFP in WT cells and in cells with the single-gene knockout of the most abundant chaperones (*Δtig, ΔdnaK*, and *ΔdnaJ*) (H) Sequences of 6 GFP mutants obtained after Sanger sequencing were aligned with the Wt GFP sequence. All the different mutations that are present in the GFP mutants are marked with a red-colored box.

**Figure S2:**
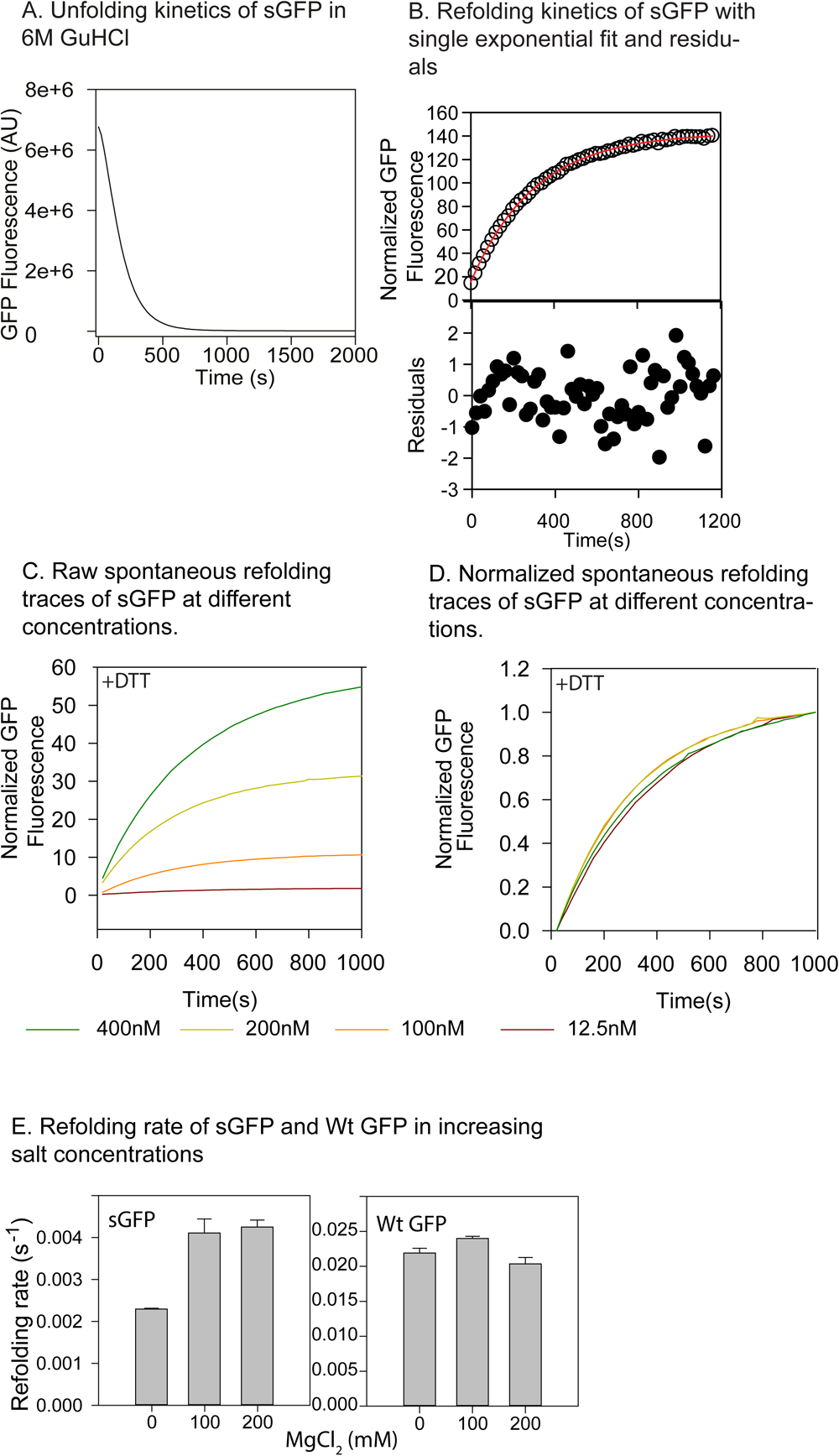
related to Figure 2. (A) sGFP was unfolded in 6 M GuHCl in buffer A for 1 hour at 25°C. Unfolding was monitored by following the decrease in fluorescence of GFP with time. (B) Refolding kinetics of spontaneously refolding sGFP in buffer A as followed by GFP fluorescence (upper panel open circles). Single exponential fit to the refolding is shown as a red line in the upper panel. The scatter from residuals are shown in the bottom panel. (C-D) sGFP refolding kinetics at different concentrations of sGFP (C). sGFP was unfolded as described earlier but at different concentrations. Refolding was initiated from each of these by a 100-fold dilution into buffer A such that the final protein concentrations were 12.5, 100, 200, or 400 nM. Refolding was monitored by measuring the recovery of GFP fluorescence over time. The plot of normalized GFP fluorescence/time (D). (E) Comparison of the refolding rate of sGFP (left panel) and Wt GFP (right panel) in the presence of different concentrations of MgCl_2_. Spontaneous refolding of both the proteins was initiated by a 100-fold dilution of the unfolded proteins in buffer A at 25°C containing different concentrations of MgCl_2_ as shown in the figure. Recovery of GFP fluorescence over time was followed to monitor refolding.

**Figure S3:**
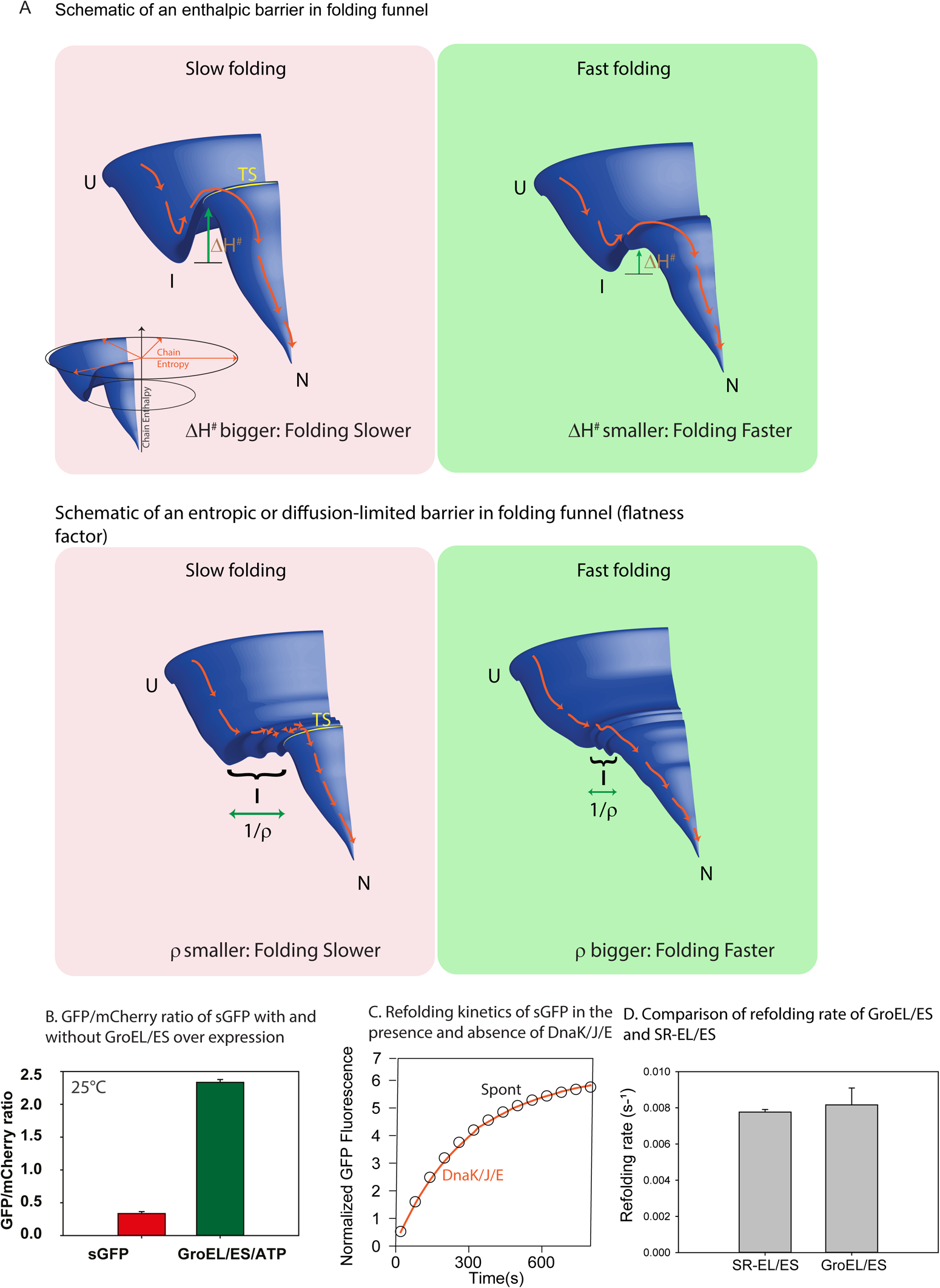
Schematic showing barriers in the folding landscape, related to Figure 2. (A) A section of the folding funnel is represented in all cases. Unfolded state (U) forms the folding intermediate (I) that are separated by a transition state (TS) from the native state (N) of the protein. Inset shows the axis - vertical axis represents enthalpy, whereas the cylindrical axis (distance from the center of rotation) represents entropy of the protein chain. The upper panel schematically shows an enthalpic barrier in folding, essentially a hill in the enthalpy axis, that leads to slower folding. A decrease in the height of the hill (decreasing ΔH^#^) leads to faster folding (right panel). The lower panel schematically depicts an entropic barrier. The intermediate (I) is a flat area in the landscape that delays folding as the conformational search is undirected and it takes time to reach TS. The flatness is inversely proportional to ρ and hence a lower ρ indicates slow folding (left panel) while an increase in ρ (decrease in the flat basin) leads to accelerated folding (right panel). (B) Bar graph showing GFP/mCherry ratio obtained from *in vivo* fluorescence of sGFP in the presence (green) and (red) absence of GroEL/ES at 25°C in the flow cytometer. (C) Refolding kinetics of sGFP in the presence and absence of DnaK/J/GrpE/ATP. Unfolded protein was diluted 100-fold in buffer A containing 800 nM of DnaK, 400 nM DnaJ, 800 nM GrpE (2:1:2), so that the final concentration of unfolded protein is 200 nM. Subsequently, refolding was initiated by adding 5 mM ATP. Recovery of GFP fluorescence over time was followed to monitor refolding. (D) Comparison of the refolding rate of sGFP in the presence of GroEL/ES and SR-EL/ES in buffer A at 25°C.

**Figure S4:**
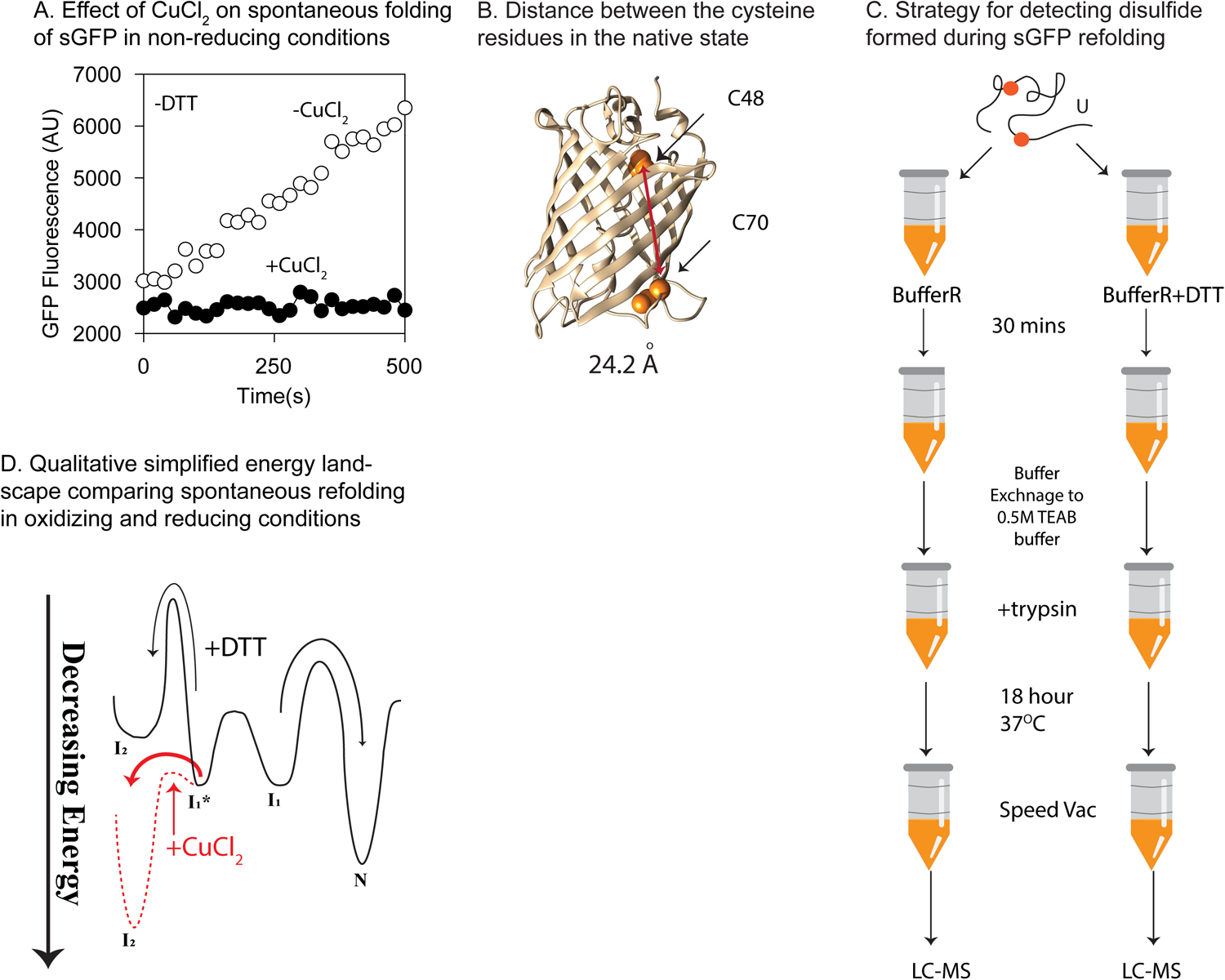
related to Figure 4. (A) Refolding kinetics of sGFP in the presence and absence of CuCl_2_. Spontaneous refolding of sGFP was initiated by a 100 fold dilution of the unfolded proteins in buffer B at 25°C. For CuCl_2_ assisted refolding unfolded protein was diluted 100-fold in buffer B containing 2 μM of CuCl_2_ so that the final concentration of unfolded protein is 200 nM. Recovery of GFP fluorescence over time was followed to monitor refolding. (B) Figure showing the distance between C48 and C70 position (Cysteine 48 and 70 (orange), connecting line between two cysteines (red)) on native GFP (using 1GFL). The figure made using Chimera. (C) Schematic showing the preparation of the sample for LC-MS. sGFP was unfolded as described earlier and refolded by 100-fold dilution in buffer A and B for 30 minutes. Buffer exchange of sGFP was done in 500 mM TEAB buffer. Then sGFP was subjected to digestion by trypsin for 18 hours at 37°C. The digested product was vacuum dried and run in LC-MS. (D) Qualitative one-dimensional energy landscape for spontaneously refolding sGFP. Black denotes the energy landscape of sGFP while refolding in the presence of DTT. The Red dashed line shows the change in the landscape in the presence of CuCl_2_. The rest of the landscape remains unchanged.

**Figure S5:**
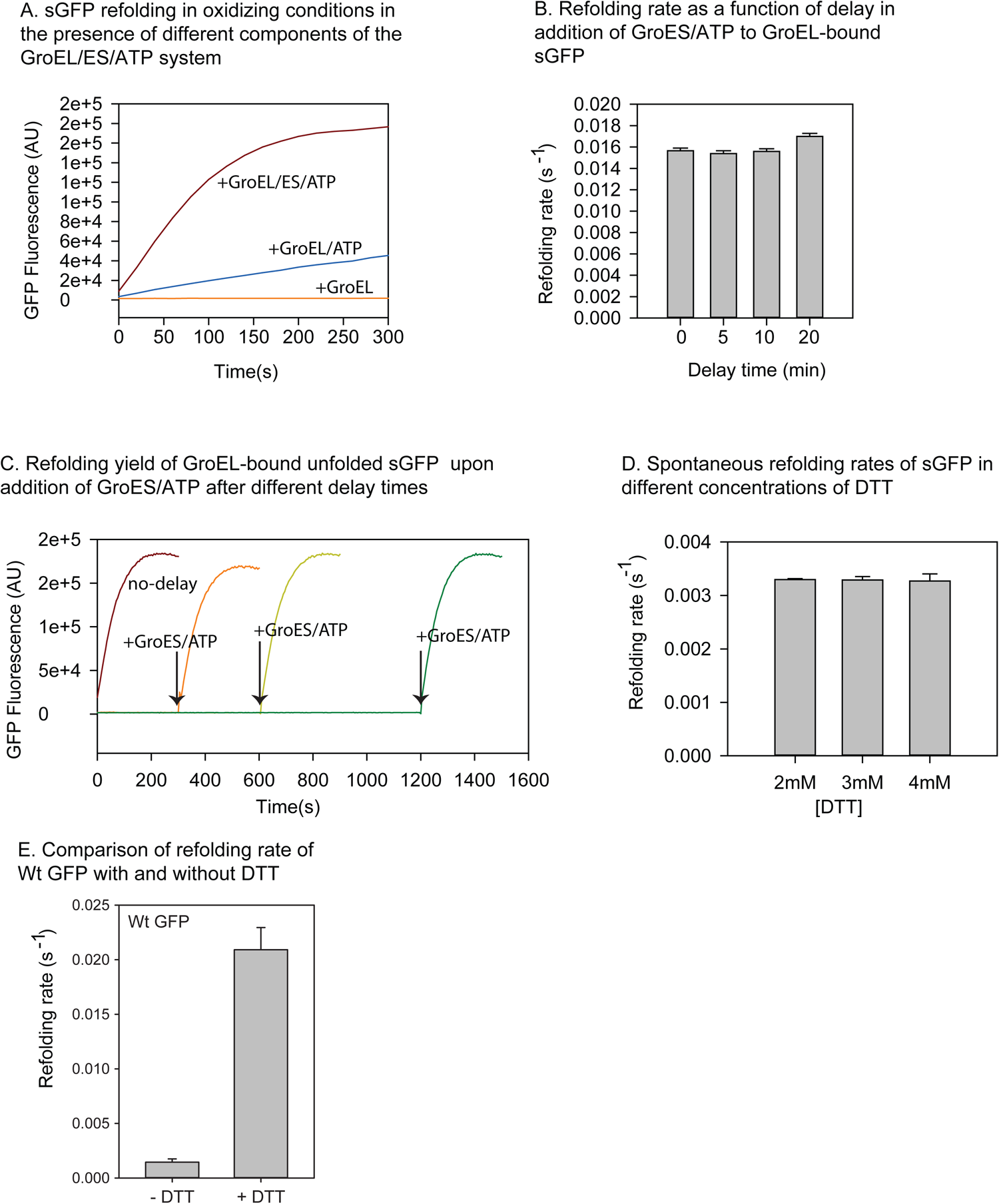
related to Figure 5. (A) Comparison of the refolding kinetics of sGFP in the presence of different components of GroEL/ES/ATP. Spontaneous refolding of sGFP was initiated as mentioned earlier in buffer B at 25°C. For the different components of GroEL/ES/ATP assisted refolding unfolded protein was diluted 100-fold in buffer C containing either GroEL alone or GroEL/ATP or GroEL/ES/ATP so that the final concentration of unfolded protein is 200 nM. Recovery of GFP fluorescence over time was followed to monitor refolding. (B) Comparison of the refolding rate of sGFP upon delayed addition of GroES/ATP to GroEL bound sGFP. Refolding of sGFP and was initiated as earlier in buffer C in the presence of GroEL, and GroES/ATP was added after a time-delay of 5, 10, or 20 minutes. For no delay, GroEL/ES/ATP was added to buffer C before adding unfolded sGFP. (C) Refolding kinetics of sGFP upon delayed addition of GroES/ATP to GroEL bound sGFP (similar to figure S5B). (D) Refolding rate as a function of DTT concentration. sGFP was unfolded as described earlier and refolding was initiated by a 100-fold dilution into buffer B containing different concentrations of DTT (2,3 and 4 mM) such that the final protein concentrations were 200 nM. Refolding was monitored by measuring the recovery of GFP fluorescence over time. Refolding traces were fitted to single exponential kinetics to obtain the refolding rates shown as bar plots with standard deviation in rates (shown as errors on the bar graph). (E) Effect of non-reducing (in buffer B, shown as - DTT) and reducing environment (in buffer A, shown as + DTT) on GFP backbone, in terms of refolding rate. Error bars indicate the standard deviation from three independent replicates.

**Table S1.** Fitted parameters from Arrhenius analysis of temperature dependent refolding reactions in different conditions as given in first column. The standard errors reported are errors obtained after fitting the triplicates of refolding rates obtained at different temperatures for each of the conditions.

| Refolding Condition | $\Delta H^\#$<br>(kCal/mole) | Std error<br>($\Delta H^\#$) | $\Delta C_p^\#$<br>(kCal/K) | Std error<br>($\Delta C_p^\#$) | $\rho$<br>(kCal/(mole.K)) | Std error<br>($\rho$) |
| --- | --- | --- | --- | --- | --- | --- |
| Spontaneous (+DTT) | -0.41 | 1.42 | 1.70 | 0.4505 | 5.88e-6 | 1.40e-5 |
| GroEL/ES/ATP | 6.20 | 0.31 | -0.18 | 0.1 | 1.26 | 0.65 |
| SREL/ES | 6.06 | 0.30 | -0.42 | 0.09 | 0.65 | 0.32 |
| wtGFP (spontaneous) | 9.26 | 0.41 | -0.24 | 0.12 | 497.30 | 339.40 |

**Table S2.**
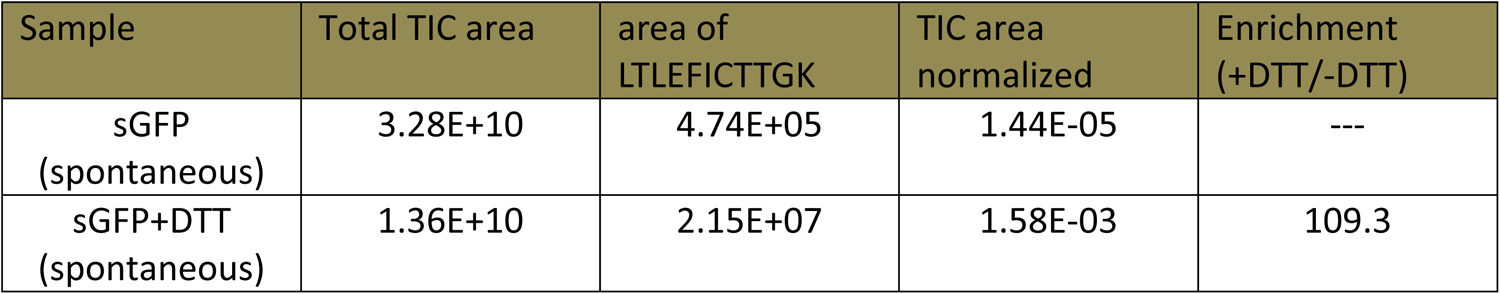
Relative quantitation and enrichment of the peptide fragment LTLEFICTTGK that is obtained from the non-disulfide bonded sGFP in refolding reactions with and without DTT. Normalization of the area of the peptide Ion count (IC) is normalized with the area under Total Ion Count (TIC) of both the LS-MS runs.

